# Spatiotemporal dynamics of signal dependent exocytosis and parasitophorous vacuolar membrane rupture during *Plasmodium falciparum* egress

**DOI:** 10.1101/2025.08.20.671297

**Authors:** Watcharatip Dedkhad, Tia Tran, Manuel A. Fierro, Carrie Brooks, Vasant Muralidharan

**Affiliations:** Center for Tropical and Emerging Global Diseases, University of Georgia, Athens, GA 30602; Department of Cellular Biology, University of Georgia, Athens, GA 30602

## Abstract

Malaria, caused by intracellular *Plasmodium* parasites, remains a major global health concern. These parasites reside and replicate within a vacuole in host red blood cells. Egress of daughter parasites out of the vacuolar and host membranes is tightly regulated via a complex mechanism. Prior studies have suggested that a cyclic-GMP driven calcium signaling pathway leads to the signal-dependent exocytosis of egress-specific vesicles that discharge several proteases into the parasitophorous vacuole. However, signal-dependent exocytosis during egress has not yet been observed in live parasitized RBCs. We targeted the exocytosis reporter, superecliptic pHlourin or SEP, to these egress-specific vesicles and utilized time-lapse imaging to observe exocytosis in live *P. falciparum* parasites. The spatiotemporal relationship between exocytosis and the breakdown of the parasitophorous vacuolar membrane (PVM) as well as parasite egress was also determined using a fluorescent reporter targeted to the PVM. Our data showed that exocytosis is triggered as early as 3 hours prior to merozoite egress. These data suggest that the PVM rupture occurs at a single site and rapidly expands from that initial site of rupture, releasing the merozoites into the RBC. This is followed by RBC membrane rupture and egress of merozoites. Using conditional mutants of the *P. falciparum* endoplasmic reticulum calcium-binding protein (PfERC), we demonstrate that knockdown of PfERC inhibits signal-dependent exocytosis of egress-specific vesicles. Together, these data demonstrate that signal-dependent exocytosis of egress-specific vesicles starts well before merozoites are formed via cytokinesis, PVM ruptures at a single site, and that PfERC is required for exocytosis of egress-specific vesicles.

## Introduction

Malaria is a life-threatening disease caused by parasites in the genus *Plasmodium*, and infection by *P. falciparum* causes the majority of the estimated mortality of nearly 600,000 in 2024 [1]. Clinical malaria manifestations of malaria result from the proliferation of *Plasmodium* parasites in human red blood cells (RBCs). *P. falciparum* parasites replicate via schizogony to produce up to 32 mature daughter parasites roughly every 48 hours. Parasite growth and replication occurs within a single membrane vacuole known as the parasitophorous vacuole (PV) within the host RBC. The PV provides a niche essential for parasite asexual development and replication by schizogony. At the end of the schizogony, the mature daughter parasites must rupture the PV membrane (PVM) and the host RBC membrane to exit or egress from the current host and invade a new RBC to continue their asexual expansion.

Egress of mature daughter parasites or merozoites is a tightly regulated complex multistep process, triggered by poorly understood signaling pathway(s) [2–11]. Egress is triggered by an unknown signal that leads to the production of cyclic GMP (cGMP), which activates Protein Kinase G (PKG) that leads to the release of intracellular calcium via unknown mechanisms [11,12]. The rise in intracellular calcium then triggers the exocytosis of egress-specific vesicles known as exonemes as well as invasion-specific vesicles known as micronemes [11,12]. Exonemes contain two key proteases, subtilisin 1 (SUB1) and plasmepsin X (PMX) that function in egress of merozoites [10,13–15]. The release of SUB1 and PMX into the PV starts a proteolytic cascade that eventually leads to breakdown of the PVM and RBC membrane [10,13–16]. However, the spatiotemporal regulation of exocytosis, PVM breakdown, and merozoite egress remains unknown.

All prior studies utilized immunofluorescence assays using fixed parasites to observe exocytosis, a dynamic process. These studies have suggested that exoneme exocytosis is upstream of PVM rupture and occurs during the final minutes before egress [8,17,18]. Our current model for *P. falciparum* egress suggests this signaling pathway occurs in the final 10 minutes of the intraerythrocytic asexual lifecycle to release merozoites from membranes [19]. This model, based on inhibitor-based studies, suggest that PKG signaling causes exocytosis of exonemes triggering a protease cascade in the PV that leads to rounding of the PVM followed by rupture of the PVM that irreversibly leads to merozoite egress [8,11,16,19]. However, since exoneme exocytosis has never been observed in live parasites, the spatiotemporal regulation of this process as well as its relationship to PVM rupture remains unknown. Furthermore, all previous studies observe microneme exocytosis, as a proxy for exoneme exocytosis, in fixed parasites [12]. However, micronemes do not function in egress and contain proteins required for merozoite invasion. This raises several questions such as: 1) When does exoneme exocytosis begin prior to merozoite egress? 2) What is the spatiotemporal relationship between exocytosis and PVM rupture? 3) What molecular mechanisms are required for signal-dependent exocytosis? To answer these questions, we wanted to directly observe exoneme exocytosis and PVM rupture in live parasites to refine our model of *Plasmodium* egress and test the role of specific proteins in these processes.

We had previously identified that the *P. falciparum* endoplasmic reticulum-resident calcium-binding protein PfERC (PlasmoDB ID: PF3D7_1108600), EF-hand calcium-binding protein, is required for merozoite egress and regulates the maturation of PMX [20]. This protein belongs to a CREC family of proteins (calumenin, reticulocalbin 1 and 3, ERC-55, and Cab) [21] and one member of this family is shown to interact directly with a protein to facilitate exocytosis [22]. Our prior studies showed that PfERC did not function in the exocytosis of micronemes, which function in parasite invasion and contain proteins required for merozoite invasion such as the Apical Membrane Antigen 1 (AMA1) [20]. However, exocytosis of exonemes, which function in egress, has not yet been observed directly. Therefore, the specific role of PfERC in exoneme egress remains unknown and we wanted to determine if PfERC functions in signal-dependent exocytosis of exonemes.

In this study, we first adapted a reporter system to observe exoneme exocytosis in live parasites during egress. The changes in PVM morphology during merozoite egress was observed using a fluorescent marker targeted to the PVM [19]. We generated parasite lines that expressed the exocytosis reporter in exonemes and a fluorescent marker on the PVM. Using these dual-labeled parasites, we observed exoneme exocytosis and PVM rupture simultaneously in *P. falciparum* infected RBCs undergoing egress. This allowed us to establish the spatiotemporal relationship between exoneme exocytosis, PVM rupture, and merozoite egress. Finally, to determine the role of PfERC in exoneme exocytosis, we generated PfERC conditional mutants expressing the exoneme-targeted exocytosis reporter. Our results show that exoneme exocytosis occurs several hours prior to egress and that PfERC is required for the signal-dependent exocytosis of exonemes.

## Results

### Targeting an exocytosis reporter to exonemes

To observe exoneme exocytosis in live parasites, we utilized a pH-sensitive variant of green fluorescent protein (GFP) known as superecliptic pHluorin or SEP [23,24] that is widely used to monitor the exocytosis and pH of subcellular compartments in several organisms including *P. falciparum* [25,26]. SEP fluorescence is quenched in acidic environments (pH <6) and is fluorescent in neutral pH environments [23,24,27,28] (Fig. 1A). Secretory vesicles are acidified as they are generated and trafficked from the Golgi. Thus, we hypothesized that SEP will not be fluorescent within acidic secretory vesicles, like exonemes, but will fluoresce when exonemes fuse to the plasma membrane and SEP is released into the neutral pH environment of the PV during exocytosis (Fig. 1A).

**Figure 1:**
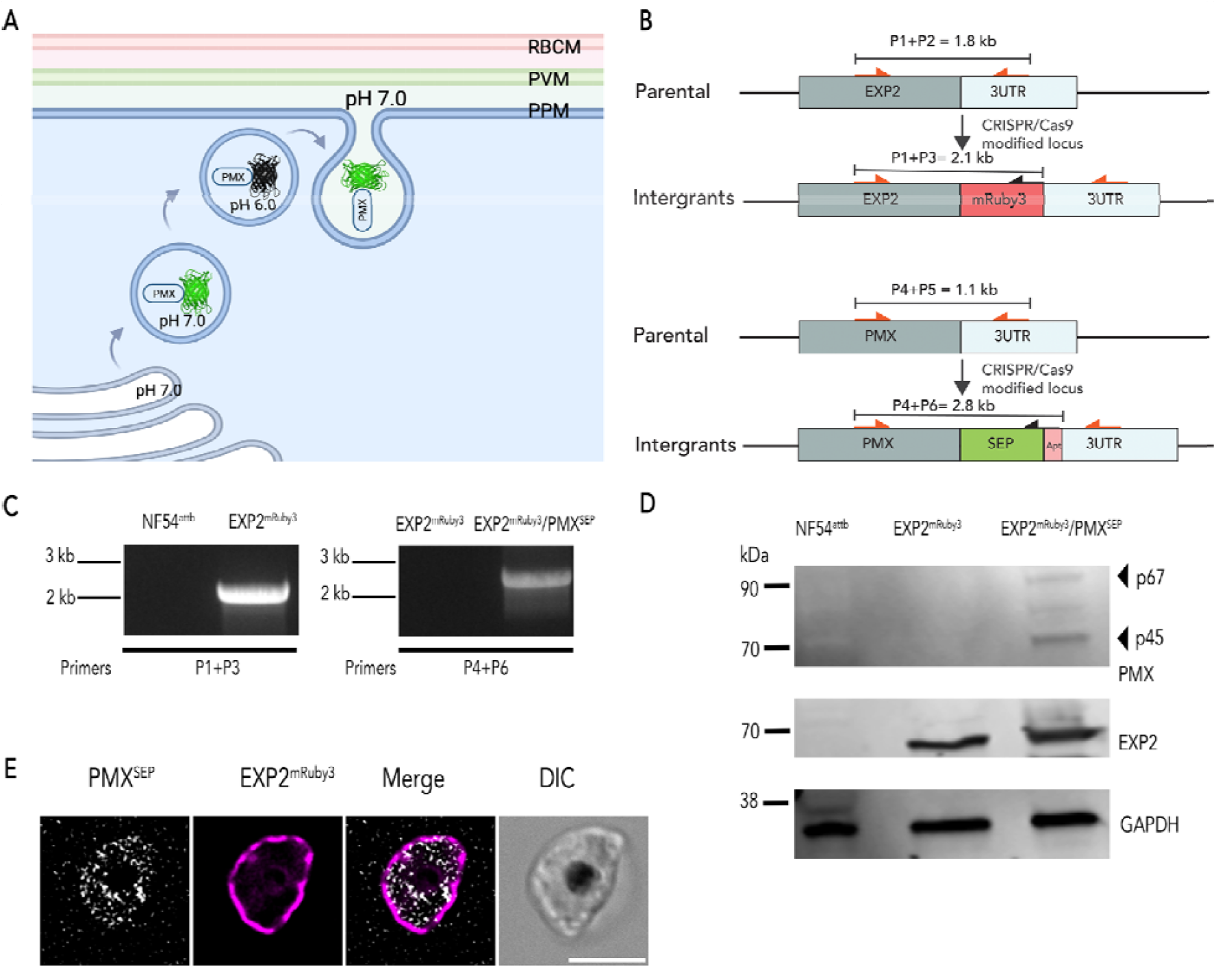
Dual-fluorescent NF54 parasites to monitor exoneme exocytosis and PVM rupture. **(A)** The Schematic shows Plasmepsin X (PMX) tagged with super-ecliptic pHluorin or SEP. SEP is non-fluorescent at pH 6.0 or less, but it is fluorescent at pH 7. **(B)** Schematics of the targeting (pMGPMX-SEP; Top) and (pbPM2-EXP2-mRuby3; bottom) plasmids, the expected recombination product, and representation of the primer pairs used to analyze an integration. **(C)** PCR diagnosis of mRuby3 fusion to the PfEXP2 locus in *P. falciparum* NF54^attB^ (left) and PMX^SEP^ in the PfEXP2^mRuby3^(right) parasites (primers sequences shown in Table S1). **(D)** Western blot of parasit lysates isolated from *P. falciparum* NF54^attB^, PfEXP2^mRuby3^, and PfEXP2^mRuby3^/PMX^SEP^ parasites probed with anti-EXP2 and anti-GFP antibodies. Results from one representative experiment of three are shown. **(E)** A representative image of live PfEXP2^mRuby3^/PMX^SEP^ schizonts (46 hpi). Images from left to right are PMX^SEP^ (gray), EXP2^mRuby3^ (magenta), fluorescence merge, and differential interference contrast or phase-contrast (DIC) Scal bar=5⍰μm.

Our current models for *P. falciparum* egress suggest that exoneme exocytosis occurs in the final minutes of egress [12,19]. To determine the relationship between exoneme exocytosis and PVM rupture, we wanted to simultaneously monitor both these processes in live parasites. Therefore, we used CRISPR/Cas9 gene editing to generate a double-labeled parasite mutant, where we tagged exported protein 2 (EXP2), a protein located in the PVM with mRuby3 (PfEXP2^mRuby3^) [29] and the exoneme protein, PMX [10,13–15], with SEP (PfEXP2^mRuby3^/PMX^SEP^; Fig. 1B, C). These data show that we successfully tagged both endogenous EXP2 and PMX gene loci with mRuby3 and SEP, respectively (Fig. 1C, D). Since PMX is essential for the completion of the intraerythrocytic lifecycle, these data also suggest that fusion of SEP to PMX did not inhibit its biological function (Fig. 1 C, D). The expression of EXP2 protein fused with mRuby3 and PMX fused with SEP was detected by western blot and showed the expected sizes, including the correct maturation species for PMX (Fig. 1D). Live imaging of late schizonts showed PMX^sep^ fluorescence as it was secreted into the PV, delineated by the EXP2^mRbuy3^ labeled PVM (Fig. 1E), indicating the tagging did not change the subcellular localization of either protein. Together, these data showed that tagging the *pmx* locus with SEP did not affect its biogenesis, maturation, function, or protein localization.

### Blocking Protein Kinase G signaling inhibits exoneme exocytosis

Using the PfEXP2^mRuby3^/PMX^SEP^ parasites, we wanted to observe the spatio-temporal dynamics of exocytosis and PVM rupture. Studies using fixed parasites in immunofluorescence assays have suggested that exocytosis of egress-specific vesicles requires PKG signaling [3,5,9,10]. Therefore, we hypothesized that exoneme exocytosis will be blocked by inhibiting PKG using specific inhibitors like compound 1 (C1) [8]. We incubated synchronized PfEXP2^mRuby3^/PMX^SEP^ schizonts at 44 hours post-invasion (h.p.i) with 1.5 µM C1. After 4 hours of incubation, the schizonts were washed and either incubated again with or without C1 and observed via live fluorescence microscopy for 30 minutes (Fig. 2).

**Figure 2:**
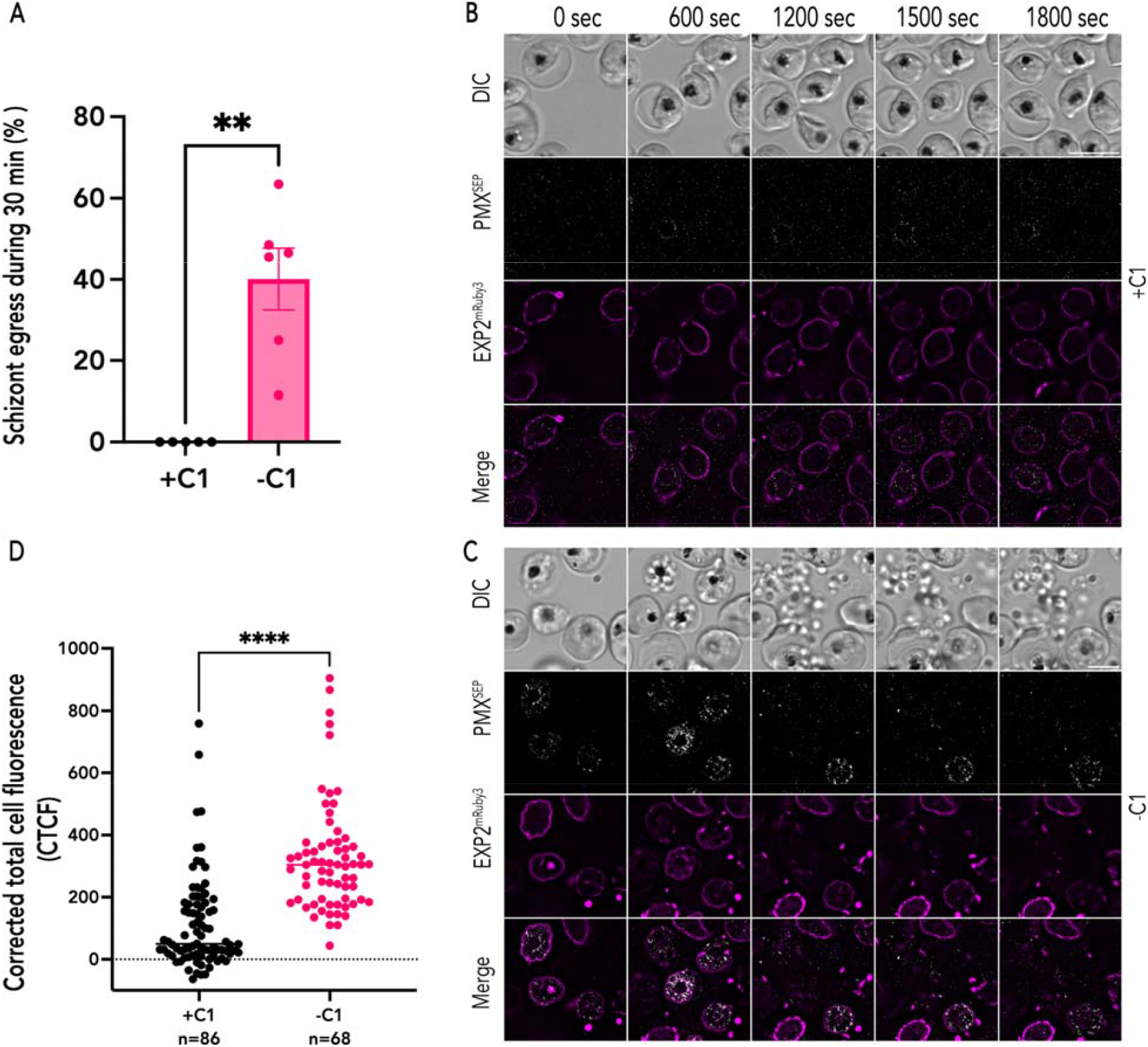
PKG inhibition blocks exoneme exocytosis. **(A)** Quantification of PfEXP2^mRuby3^/PMX^SEP^ parasite egress in the presence of C1 (+C1; black) or absence of C1 (−C1; magenta) during the 1800-second recording period. Each dot represents data from one time-lapse imaging experiment (n = 5 recordings from 3 biological replicates for PfEXP2^mRuby3^/PMX^SEP^ parasites with C1, n= 6 recordings from 3 biological replicates for PfEXP2^mRuby3^/PMX^SEP^ parasites without C1, error bars = SEM; **p < 0.001 by unpaired two-tailed t test) **(B)** A representative image from live imaging of synchronized PfEXP2^mRuby3^/PMX^SEP^ schizonts in the presence of C1. **(C)** A representative imag from live imaging of synchronized PfEXP2^mRuby3^/PMX^SEP^ schizonts after C1 removal. Parasite egress occurs, and free-merozoites are scattered in the extracellular space (DIC). **(D)** Corrected total cell fluorescence (CTCF) of PMX^SEP^ quantified from time-lapse images of synchronized PfEXP2^mRuby3^/PMX^SEP^ schizonts incubated with (black; n=86, 3 biological replicates) or without (magenta; n=68, 4 biological replicates) C1. ***p-value<0.0001, unpaired t-test. error bars = SEM. Scale bar=5⍰μm.

We observed that in the presence of C1, PfEXP2^mRuby3^/PMX^SEP^ schizonts did not egress during the 30 minute window of observation (Fig. 2A and Movie S1). The PVM was intact in these schizonts during this time period (Fig. 2B). These data show that blocking exocytosis via PKG inhibition also inhibits PMX^SEP^ fluorescence (Fig. 2B), indicating that the environment within unsecreted exonemes is acidic resulting in quenching of SEP fluorescence (Fig. 2B). In some schizonts incubated with C1, we observed SEP fluorescence, which may be due to exocytosis of exonemes prior to C1 addition (Fig. 2). However, none of these schizonts were able to rupture the PVM or egress during the observation period (Fig. 2A).

In the absence of C1, we observed punctate PMX^SEP^ fluorescence within the PVM, indicating PMX^SEP^ was exposed to neutral pH and suggesting that exoneme exocytosis had occurred (Fig. 2C and Movie S2). The time-lapse images are snapshots at a single timepoint and as the vesicle pH is neutralized upon fusion to the parasite plasma membrane a punctate PMX^SEP^ fluorescent signal was observed from the neutralized exoneme (Fig. 2C and Movie S2). This is followed by rapid dissolution of the fluorescence due to the rapid dilution of PMX^SEP^ into the much larger PV compartment. These data suggest that exoneme exocytosis occurs all over the schizont and may not be limited to a single exit site or specific cellular location within the merozoite. In the absence of C1, exoneme exocytosis occurred first, then the PVM ruptured in most PfEXP2^mRuby3^/PMX^SEP^ schizonts followed by the release of free merozoites (Fig. 2C). The corrected total cell fluorescence for each schizont observed via live video microscopy was integrated across several time points (Fig. 2D). These data show that in the presence of C1, PMX^SEP^ fluorescence signal is substantially reduced suggesting that exocytosis does not occur when PKG is inhibited (Fig. 2D).

One possibility we considered is that the reduction in PMX^SEP^ fluorescence signal is a result of direct interference from C1. Thus, we utilized another PKG inhibitor, ML10, that also inhibits PKG but is chemically distinct from C1 [30]. Incubating PfEXP2^mRuby3^/PMX^SEP^ schizonts with ML10 also resulted in loss of PMX^SEP^ fluorescence, suggesting that PMX^SEP^ is in an acidic vesicle when exocytosis is blocked (Supplementary Fig. 1). Thus, these data show that exonemes are acidic secretory vesicles, and that their exocytosis is dependent upon PKG signaling.

### Spatiotemporal relationship between exoneme exocytosis and PVM rupture

The spatio-temporal relationship between exoneme exocytosis and merozoite egress has been determined based on studies that utilized fixed parasites [12]. More recently, live imaging approaches have been utilized to investigate the morphology of the PVM during egress [11,19]. These data show that a calcium-dependent signal causes the rupture of the PVM followed by RBC membrane breakdown to release the merozoites [11,19]. However, a central part of the egress model, the signal-dependent exocytosis of exonemes has not yet been observed by live microscopy. Therefore, we wanted to define the relationship between signal-dependent exocytosis of exonemes and PVM rupture.

To determine the relationship between signal-dependent exocytosis of exonemes and PVM rupture, synchronized PfEXP2^mRuby3^/PMX^SEP^ schizonts (44 h.p.i) were prepared as described previously and incubated with C1 for 4 hours. After removing C1, we observed the schizonts via time-lapse imaging for 30 mins with 30 sec intervals (Fig. 3 and Movie S3). As others have reported [11,19], we observed irregularly shaped PVM at the start of our observation (Fig. 3A). About 8 minutes prior to egress, the PVM adopted a rounded shape (Fig. 3) and this was followed rapidly (~1 minute) by the first observable break in the PVM (Fig. 3B). The schizonts egressed about 7 minutes after the PVM rupture (Fig. 3C). Our observations agree with prior studies that also utilized reversible PKG inhibitors to establish the timeline of egress starting from PVM rounding [11,19]. Thus, despite differences in parasite lines and preparation across labs, as well as microscopy settings, the observed timeline of the egress cascade is consistent.

**Figure 3:**
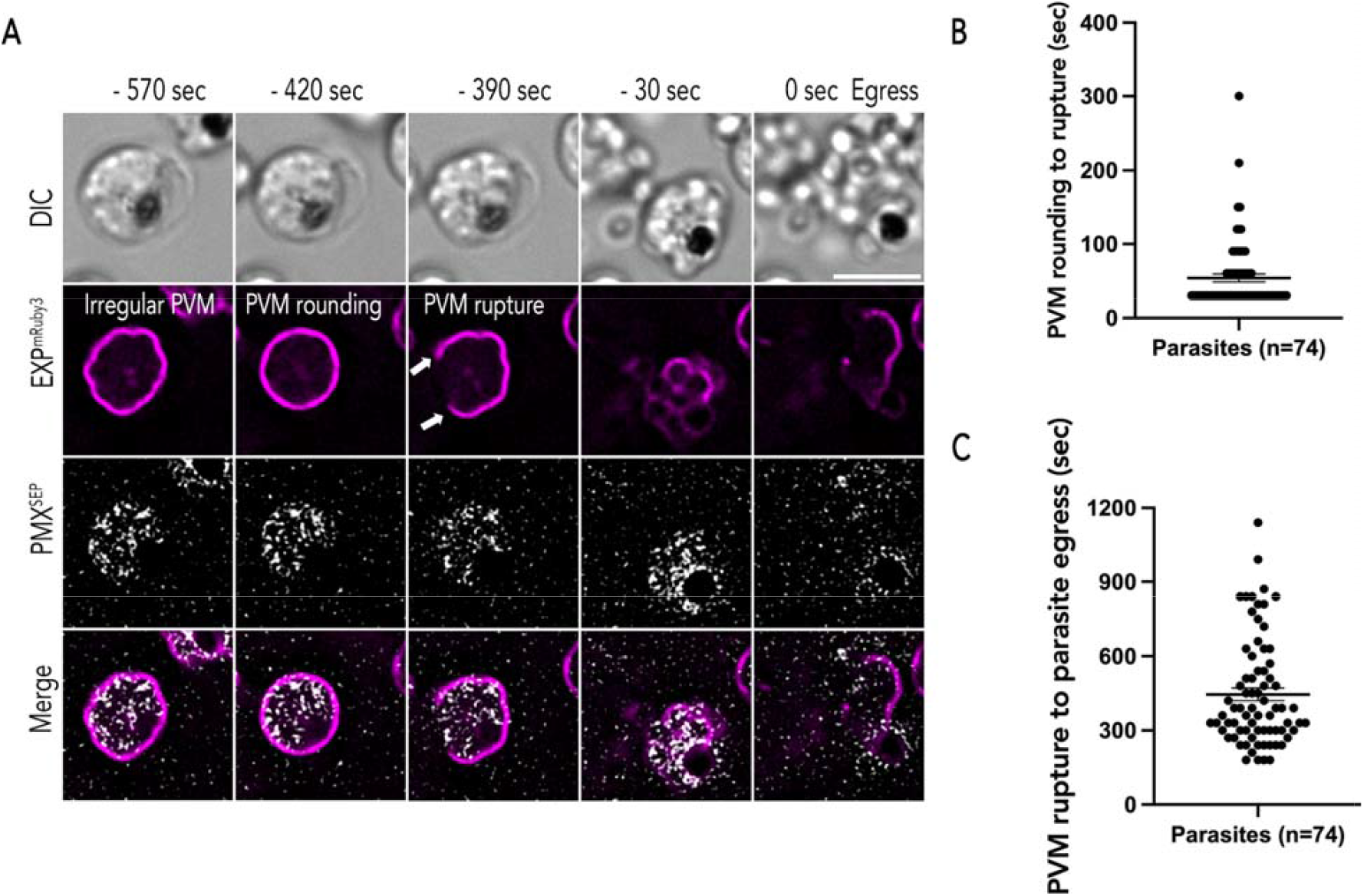
Exoneme exocytosis prior to egress. **(A)** Representative time lapse images (captured at 30 sec intervals) during egress of PfEXP2^mRuby3^/PMX^SEP^ merozoites. Minus time indicates time points before merozoite egress. PVM (magenta) morphology changes from an irregular shape to a rounded shape. In most schizonts, PVM rupture (white arrows) occurred in the next frame or two (30 to 60 sec) after PVM rounding. Merozoites free from membranes indicate egress. Several exoneme exocytosis (gray) events were observed from the start of the recording until merozoite egress. Scale bar=5 μm. **(B)** The estimated time between PVM rounding to rupture (Mean ± SEM = 53.919 ± 5.321 sec) and **(C)** the time between PVM rupture to merozoite egress (Mean ± SEM = 445.946 ± 25.276 sec). n = 74, 7 biological replicates, double-labeled parasites, error bars = SEM, time-lapse recordings of parasite egress is a 30-sec interval.

Since our images were taken using 30 sec imaging intervals, we wanted to observe the exact temporal relationship between PVM rounding and PVM rupture as well as observe PVM rupture at a higher temporal resolution. Therefore, we utilized 4 sec and 2 sec time-lapse imaging to observe egressing PfEXP2^mRuby3^/PMX^SEP^ schizonts (Fig. 4). Similar to our prior observations (Fig. 3), we observed that the PVM morphology changed from an irregular shape to a more rounded shape (Fig. 4A, B, Supplementary Fig. 2A-C and Movie S4, S5). The diameter (~4.3 μm) and circumference (~13.6 μm) of the PVM in all rounded schizonts were similar across biological replicates (Fig. 4B,C).

**Figure 4:**
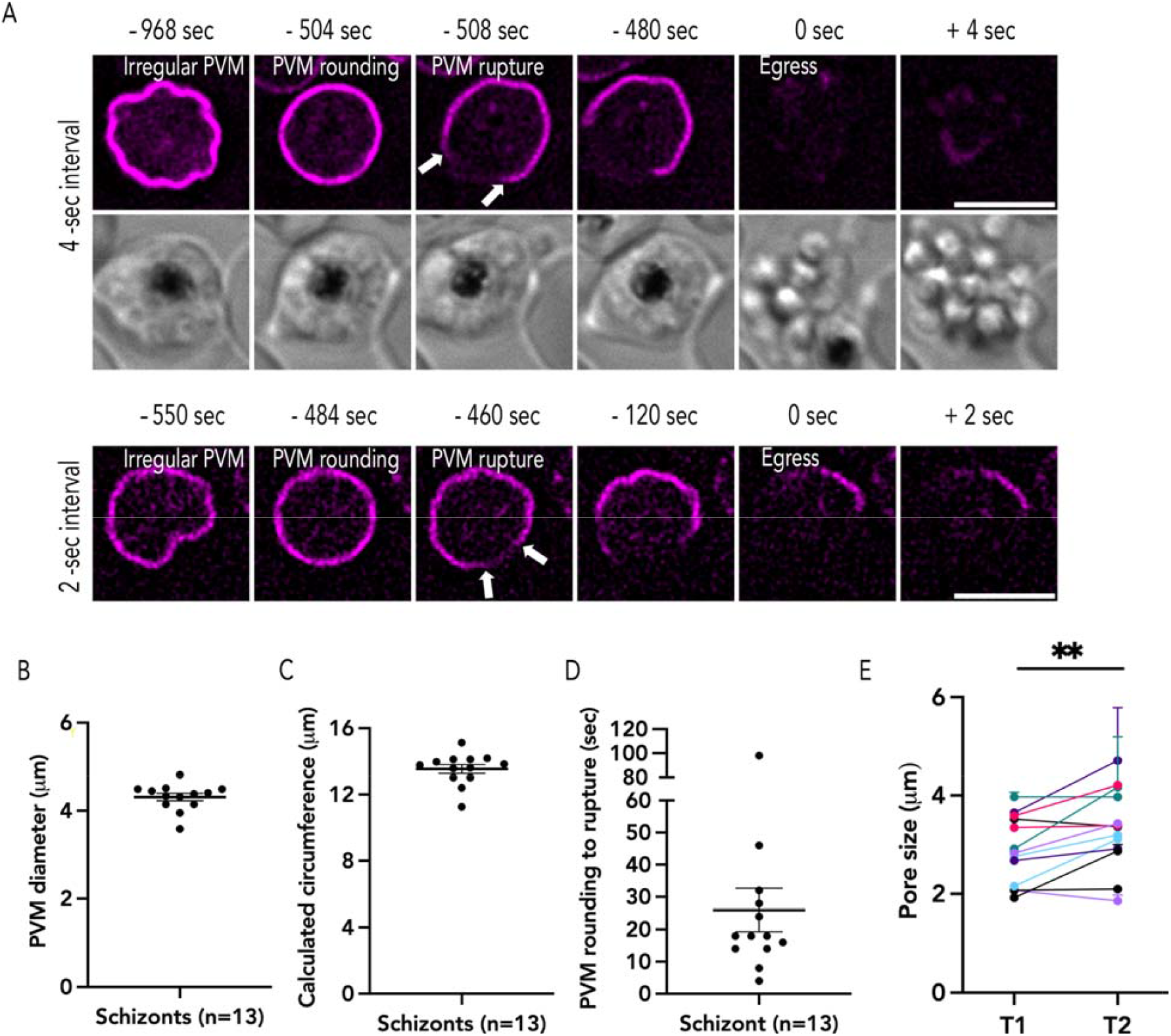
PVM rupture occurs at a single site. (A) Representative time lapse images captured from 4-sec (top) and 2-sec intervals (bottom), the white arrow indicates two ruptured ends of the PVM **(B)** The diameter of rounded PVM from 2-sec interval time lapse images (Mean ± SEM= 4.31 ± 0.08 μm), **(C)** circumference (Mean ± SEM = 13.54 ± 0.264 μm), **(D)** the duration of PVM rounding (Mean ± SEM = 26 ± 6.706 sec; n= 13, 4 biological replicates). (E) The pore size changes between the first time point PVM breakdown (T1), Mean ± SEM = 2.88 ± 0.191 μm, and the second time point (T2), Mean ± SEM =3.33 ± 0.226 μm (each color represents one schizont imaged every 2 sec, n=13, 4 biological replicates, error bars = SEM, paired t-test with ** p ≤ 0.001). The measurement data are shown in Supplementary Fig. 2. Scale bar = 5⍰μm.

Within a few seconds (~25 sec) after the schizont rounds, the PVM ruptures (Fig. 4D), though we observed some variation in the time taken for PVM rupture after rounding (Fig. 4D). In some schizonts, it took nearly a minute to observe PVM rupture after rounding, while in other schizonts PVM rupture occurred a few seconds (<10 sec) post-rounding (Fig. 4D). These data support the hypothesis that PVM rounding is a critical morphological step for PVM rupture to occur (Fig. 4D; [11,19]. Regardless of the time interval of imaging, PVM rupture always occurred at a single site after rounding (Fig. 4). The size of this initial rupture was consistent across several biological replicates (~2.8 μm; Fig. 4E, Supplementary Fig. 2 D-E). We never observed a second site of PVM rupture during egress and the initial rupture widened in subsequent time-lapse images allowing the merozoites to escape the PVM (Fig. 4E). These data suggest that the PVM rupture may be regulated and likely occurs at a specific site on the membrane (Fig. 4).

### Exoneme exocytosis begins several hours prior to merozoite egress

Our studies using reversible PKG inhibitors showed that exoneme exocytosis was already occurring in PfEXP2^mRuby3^/PMX^SEP^ schizonts at the start of our observation (Fig. 2-4). Therefore, we wanted to establish a time-line for signal-dependent exoneme exocytosis and study this process without the use of any inhibitors. Using highly synchronized PfEXP2^mRuby3^/PMX^SEP^ parasites (44 h.p.i), we monitored exoneme exocytosis until merozoites egressed from these schizonts (Fig. 5 and Movie S6).

**Figure 5:**
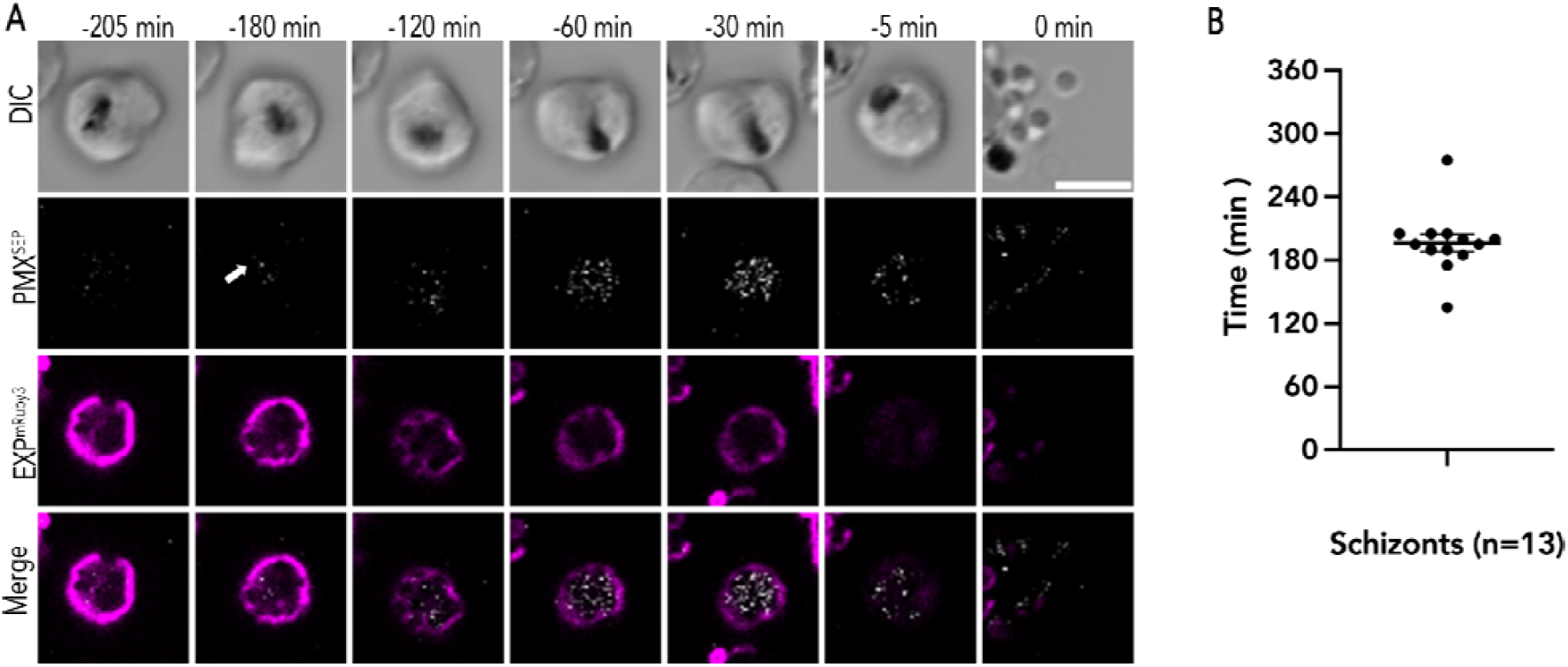
Exoneme exocytosis begins 3 hours prior to egress. **(A)** Representative time lapse images of PfEXP2^mRuby3^/PMX^SEP^ merozoite exocytosis and egress. The arrow indicates the first image where fluorescence signals were observed. Time points prior to merozoite egress (0 min) are indicated. Scale bar=5⍰μm. **(B)** The average time of exoneme exocytosis observed in PfEXP2^mRuby3^/PMX^SEP^ schizonts prior to egress. (n= 13 schizonts from 4 biological replicates, Mean ± SEM = 196.53 ± 8.35 min).

We observed several naturally egressing PfEXP2^mRuby3^/PMX^SEP^ schizonts over four biological replicates and at the start of the recording we did not observe any PMX^SEP^ fluorescence in any of these schizonts indicating that exoneme exocytosis had not yet started (Fig. 5). Parasite viability was assessed using the movement of schizonts within the infected RBC as well as by hemozoin movement within the food vacuole and only viable schizonts that egressed over this time period were analyzed further (Fig. 5). Moreover, the changes in PVM morphology as assessed by PfEXP2^mRuby3^ fluorescence was another measure of parasite viability over the observation time period (Fig. 5A and Movie S6).

In all PfEXP2^mRuby3^/PMX^SEP^ schizonts, we observed punctate PMX^SEP^ fluorescence within the PVM starting several hours (~3 hours) prior to merozoite egress (Fig. 5B). Once the punctate PMX^SEP^ fluorescence started, these events were observed continuously until the merozoites egressed from the infected RBC (Fig. 5A and Movie S6). These data suggest that the signals that kickstart exoneme exocytosis occur at least 3 hours prior to egress (Fig. 5). Another possibility is that the observed PMX^SEP^ fluorescence at this earlier timepoint could be secretory vesicles that were not yet acidified and therefore fluorescent. The process of vesicle acidification is intrinsic to the secretory pathway and independent of PKG signaling. In this case, PKG inhibitors would not inhibit PMX^SEP^ fluorescence within vesicles that were not yet acidified. Therefore, to distinguish between these possibilities, we incubated similarly isolated PfEXP2^mRuby3^/PMX^SEP^ schizonts (44 h.p.i) with the PKG inhibitor ML10 over the same time period (Supplementary Fig. 3 and Movie S7). We did not observe any PMX^SEP^ fluorescence indicating that SEP was already present in the acidic exonemes and that the observed fluorescence was not due to immature secretory vesicles that were not yet acidified (Supplementary Fig. 3 and Movie S7). Together, these data suggest that a critical concentration of egress proteases may be needed in the PV to activate merozoite egress, which may take several hours to achieve. Alternatively, there may be a second signal that activates the egress protease cascade and PVM rounding [11,19].

A recent study showed that *P. falciparum* V-type ATPases may localize to secretory vesicles in merozoites [31] and therefore, we wanted to determine if these V-type ATPases function to maintain the acidic pH within exonemes. To test this, we incubated PfEXP2^mRuby3^/PMX^SEP^ late schizont with ML10 to prevent exocytosis as well as with the V-type ATPase-specific inhibitor Bafilomycin A1 (BaF). We hypothesized that incubation with ML10 will prevent changes in PMX^SEP^ fluorescence due to exocytosis and the addition of BaF will inhibit the V-type ATPase leading to neutralization of the vesicle pH that will result in PMX^SEP^ fluorescence (Supplementary Fig. 4). No exocytosis or egress was observed in ML10-treated PfEXP2^mRuby3^/PMX^SEP^ parasites (Supplementary Fig. 4A). Surprisingly, we did not see an increasing fluorescence during the 15-min time-lapse videos in ML10-treated PfEXP2^mRuby3^/PMX^SEP^ parasites incubated with BaF (Supplementary Fig. 4B). There was no discernible difference in PMX^SEP^ fluorescence upon incubation with BaF (Supplementary Fig. 4B). These data suggest that the mechanisms maintaining the acidic pH within exonemes are insensitive to BaF (Supplementary Fig. 4). The data suggest there may be other BaF-insensitive proton pumps or proton exchangers that may play a role in maintaining the pH within exonemes.

### PfERC is required for exoneme exocytosis

Calcium signals are known to be essential for exoneme exocytosis but the mediators of this calcium signal within the secretory pathway have not yet been identified [12,19]. We had previously identified an ER-resident calcium binding protein, PfERC, that was essential for merozoite egress [20]. Our data showed that PfERC functions in the egress proteolytic cascade but is not required for ER calcium homeostasis or for microneme exocytosis [20]. Since egress proteases are stored within exonemes and released via exocytosis into the PV during egress, interfering with either exoneme biogenesis or exocytosis may result in loss of protease activity or function. Therefore, we hypothesized that PfERC is required for egress protease function because it may function in exoneme biogenesis or exocytosis.

To test this hypothesis, we first generated PfERC conditional mutants using the glucosamine (GlcN) responsive ribozyme system (Fig. 6A) [20,32]. The addition of GlcN activates the *glmS* but not *M9* ribozyme, resulting in cleavage of mRNA that leads to mRNA degradation and protein knockdown (Fig. 6B). The endogenous locus of PfERC in the NF54 *P. falciparum* parental strain was fused to three copies of the hemagglutinin (HA) tag and either the inducible *glmS* (termed PfERC-*glmS*) ribozyme or the inactive *M9* (termed PfERC-*M9*) ribozyme (Fig. 6A) [20,32]. Clonal PfERC-*glmS* and PfERC-*M9* parasite lines were generated from independent transfections (Supplementary Fig. 5A, B). The PMX locus in these clonal PfERC-*glmS* and PfERC-*M9* parasite lines was then tagged with SEP to generate double mutants (termed PfERC-*glmS*/PMX^SEP^ and PfERC-*M9*/PMX^SEP^; Supplementary Fig. 5C, D). In these PfERC-*glmS*/PMX^SEP^ and PfERC-*M9*/PMX^SEP^ double mutants, PMX expression was also regulated by the *tetR* aptamer system [20,33] (Supplementary Fig. 5).

**Figure 6:**
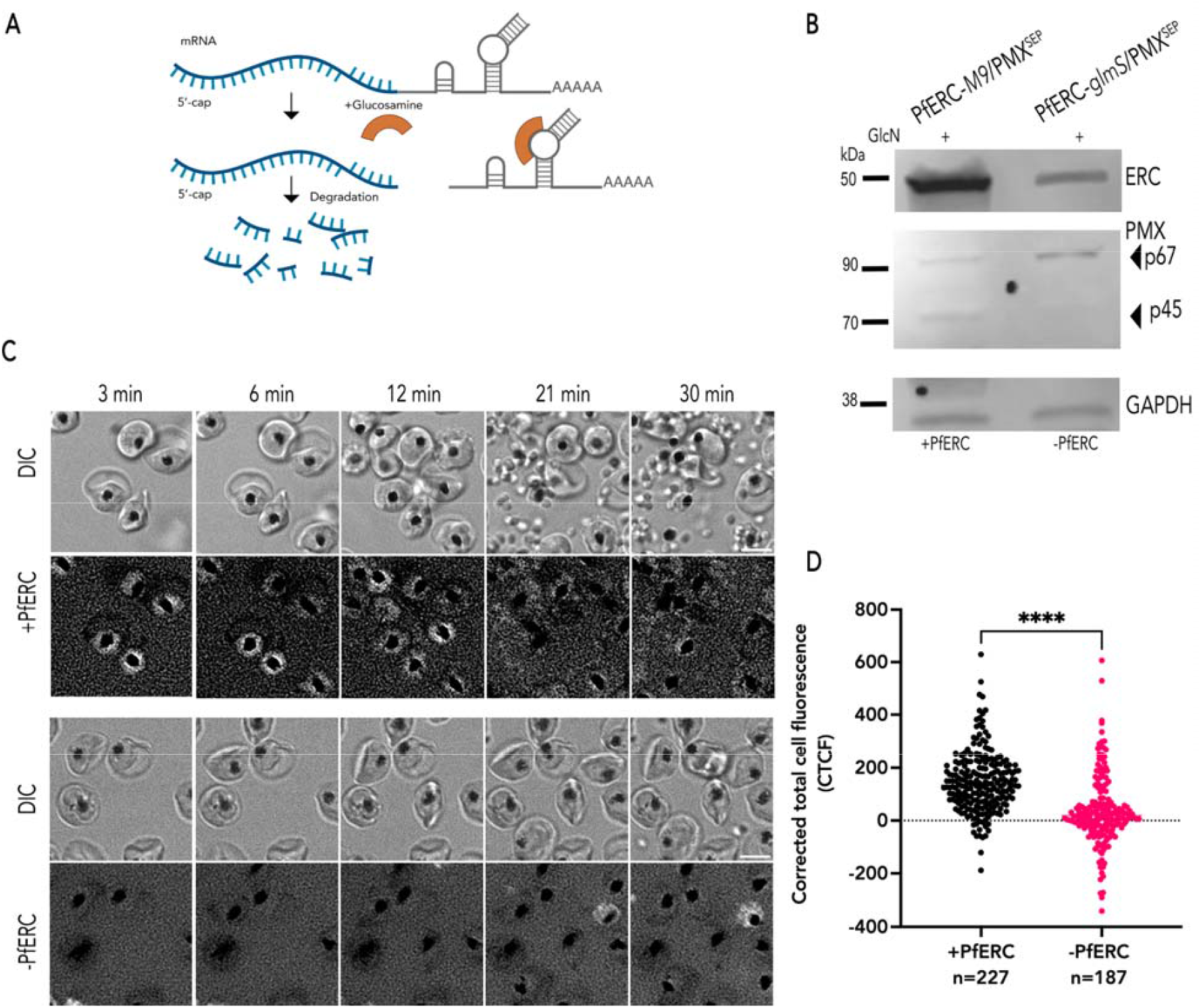
PfERC is required for exoneme exocytosis in live parasites (A) Schematic shows the mechanism of The condition that knocks down the *glmS* ribozyme system. *glmS* is an inactive ribozyme that is transcribed but not translated with the mRNA of a protein of interest. Adding glucosamine (GlcN) leads to phosphorylation within The cell to glucosamine-6-phosphate (GlcN-6P). GlcN-6P binds to the transcribed mRNA, and the *glmS* ribozyme is activated and cleaves itself from the mRNA. This leads to disassociation of the mRNA from its poly(A) tail and results in the degradation of target-specific mRNA. The resulting decline in mRNA levels leads to reduced protein levels and, thus to loss of gene expression. As a control, we generated parasite lines containing a mutated version of the *glmS* ribozyme, called M9, which cannot cleave itself upon binding of GlcN. (B) Representative of western blot of parasite lysates isolated from PfERC-*glmS*/PMX^SEP^ and PfERC-*M9*/PMX^SEP^ schizonts, PfERC-*glmS* schizonts grown in the presence of GlcN for 48⍰h and probed with anti-HA antibodies (top panel), anti-GFP antibodies (middle panel), and anti-GAPDH (loading control; bottom panel). (C) Representative images from Glc-treated PfERC-*glmS*/PMX^SEP^ and PfERC-*M9*/PMX^SEP^ schizonts from live fluorescence microscopy. Phase contrast (top) and fluorescence (bottom) images. (D) Corrected total cell fluorescence (CTCF) integrated over the entire video (n= 10 recordings from 3 biological replicates for Glc-treated PfERC-*M9*/PMX^SEP^, n= 11 recordings from 3 biological replicates for Glc-treated PfERC-*glmS*/PMX^SEP^, ****p-value<0.0001,unpaired t-test). Scale bar=51μm

Parasites were prepared as previously to obtain highly synchronized PfERC-*glmS*/PMX^SEP^ and PfERC-*M9*/PMX^SEP^ ring stage parasites (2 h.p.i). The 2 h.p.i ring stage parasites were incubated with 7.5 mM GlcN for 48 h. The addition of GlcN led to a reduction in PfERC expression in PfERC-*glmS*/PMX^SEP^ schizonts, while there was no reduction in PfERC expression in PfERC-*M9*/PMX^SEP^ schizonts grown under identical conditions (Fig. 6B). These data confirmed that in these double mutants the addition of GlcN led to knockdown of PfERC only in PfERC-*glmS*/PMX^SEP^ parasites (termed −PfERC) but had no effect on PfERC expression in PfERC-*M9*/PMX^SEP^ parasites (termed +PfERC; Fig. 6B). Our prior work had shown that PfERC knockdown led to a block in PMX activity [20,33]. The addition of GlcN to PfERC-*glmS*/PMX^SEP^ and PfERC-*M9*/PMX^SEP^ parasites also resulted in loss of PMX processing (Fig. 6B).

Next we wanted to test if PfERC knockdown had any effect on exoneme exocytosis. As before, we treated ring stage (2 h.p.i) PfERC-*glmS*/PMX^SEP^ and PfERC-*M9*/PMX^SEP^ parasites with GlcN and then these GlcN treated schizonts (44 h.p.i) were incubated with C1 for 4 hours to prevent egress, allow schizonts to fully develop and become egress competent. At 48 h.p.i, C1 was washed off from these schizonts and we observed these parasites using wide-field live fluorescence microscopy (Fig. 6C). Time-lapse images were collected every 45 seconds for 30 minutes (Fig. 6C). During the 30-min video, GlcN-treated PfERC-*MP*/PMX^SEP^ schizonts were able to egress (Fig. 6C and Movie S8). Several fluorescent events were observed in PfERC-*M9*/PMX^SEP^, while almost no fluorescence events were observed in GlcN-treated PfERC-*glmS*/PMX^SEP^ (Fig. 6C and Movie S9). The corrected total cell fluorescence for each observed schizont within the field was integrated over 30 minutes (Fig. 6D). These data show that incubating PfERC-*M9*/PMX^SEP^ with GlcN does not have an effect on exoneme exocytosis or egress (Fig. 6D). In contrast, incubating PfERC-*glmS*/PMX^SEP^ with GlcN results in loss of exoneme secretion and blocks egress (Fig. 6D). These data show that PfERC is required for exoneme exocytosis (Fig. 6C, D).

## Discussion

Merozoite egress from the infected RBC is a synchronized multistep process that is regulated in a timely manner [12]. Our current model for parasite egress suggests that two second messengers, calcium and cGMP, are critical for *Plasmodium* egress [12]. The molecular mechanisms linking calcium and cGMP are still unknown, but both these second messenger pathways are thought to converge on the exocytosis of egress-specific vesicles known as exonemes. Exoneme exocytosis is essential for parasite egress as it results in the release of two well-characterized proteases into the PV, SUB1 and PMX [10,13–16]. The release of SUB1 and PMX into the PV eventually results in PVM rupture, leading to parasite egress [12]. However, the signal-dependent exocytosis of exonemes has not yet been observed in live *Plasmodium* schizonts during parasite egress. Thus, we aimed to determine the spatio-temporal relationship between the release of PMX into the PV and eventual rupture of the PVM. In this study, we used multiple approaches, including genetic tools combined with live fluorescence microscopy, to investigate two critical steps during egress: exoneme exocytosis and PVM rupture in live parasites. Based on our results, we propose a timeline of *Plasmodium* egress from the infected RBC (Fig.7).

Studying exocytosis in live *Plasmodium* poses many challenges. Merozoites are only 1 μm in size, and nearly two dozen merozoites are packed in the PVM inside the ~7 μm diameter of the host RBC. Moreover, a group of merozoites are moving around the residual body within the schizont, which makes it extremely challenging to observe individual merozoites as well as to track the path of secretory organelles rapidly undergoing exocytosis. A reporter that provides a signal only upon exocytosis is ideal to monitor this process within *P. falciparum* schizonts. Therefore, in this study we used pH-sensitive variant of green fluorescent protein (GFP) known as superecliptic pHluorin or SEP [23,24] to monitor the exocytosis of exonemes during *Plasmodium* egress together with PVM morphology as well as determine if PfERC functions in this process. We targeted SEP to exonemes by fusing it with PMX, which did not interfere with PMX processing or function. Our data show that PMX^SEP^ is non-fluorescent until exocytosis, strongly suggesting that exonemes are acidic secretory vesicles. Surprisingly, we did not observe loss of acidification when we used the V-type ATPase proton pump inhibitor BaF. A prior study localized the *P. falciparum* V-type ATPase to secretory vesicles in schizonts and showed that BaF leads to acidification within the parasite [31]. However, in this previous study [31], addition of BaF to late schizonts did not inhibit parasite egress. Since the proteases essential for egress are activated by low pH[15,34–37], this suggests that the V-type ATPase may not be responsible for acidification of exonemes. Proton pumping pyrophosphatases are another mechanism to acidify intracellular organelles and *P. falciparum* encodes for two pyrophosphatases, PfVP1 (PF3D7_1456800) and PfVP2 (PF3D7_1235200). PfVP1 localizes to the parasite plasma membrane and is required for ring to trophozoite stage transition [38] while PfVP2 is not essential for the intraerythrocytic life cycle [29,39]. Thus, it is not known how exonemes are acidified and more research is needed to identify the mechanism responsible for exoneme acidification.

PMX^SEP^ is non-fluorescent within the acidic exonemes and we observe PMX^SEP^ fluorescence when it is exposed to the neutral environment within the PV upon signal-dependent exocytosis. We detected exocytosis signals as a punctate pattern inside the enclosing PVM. There was no reproducible change observed in the frequency or intensity of exocytosis during egress. Given that merozoites moved around the residual body, we could not track the path of each exocytosis event. Due to the resolution limits of microscopy, it was also not possible to locate the exact site of exoneme exocytosis or if it occurred at the apical end of merozoites. Nevertheless, our data show that blocking PKG signaling inhibits PMX^SEP^ fluorescence. Studies utilizing reversible inhibitors suggest that the cGMP and calcium signaling pathways that regulate merozoite egress are activated in the final minutes of the intraerythrocytic life cycle [2–11,19]. Our data suggest that exoneme exocytosis begins several hours prior to egress and continues until parasite egress. Since this process is dependent upon PKG signaling, these data further suggest that the cGMP signaling pathway may be activated several hours before merozoite segmentation is completed. These data are in agreement with studies showing that blocking merozoite segmentation does not inhibit parasite egress [40]. This is likely because, as our data shows, the egress pathway starts before segmentation begins. It is also possible that the cGMP and calcium signals that kickstart exoneme exocytosis about 3 hours prior to egress are similar though not identical to the calcium signaling pathway that results in PVM rounding [2–11].

Even though our data show that the egress proteases are released into the PV several hours prior to egress, it is not clear whether they are active in the PV during this time before egress. The egress proteases are activated by acidic pH [15,34–37] and it is possible that in the neutral pH of the PV, the proteases need several hours to ensure complete processing of their substrates. Another possibility is that they may have specific interacting partners that lead to an enhancement of protease activity [41]. More studies are needed to understand how the egress proteases function within the PV to finish the processing of their substrates to ensure the merozoites are invasion competent prior to egress.

Our data are in agreement with prior studies that show that PVM rounding occurs about 8-10 minutes prior to egress [2–11,19]. A minute after PVM rounding, the PVM ruptures and data show that the timescale for rupture after membrane rounding was similar regardless of imaging timescale. The imaging data further show that the PVM ruptures with a <3 μm pore at a single site on the membrane, which then rapidly expands. In all our biological replicates, we always observed only one site of PVM rupture. The current model for egress suggests that PVM poration occurs because exonemes secrete proteases into the PV minutes prior to egress [8,10,13–18]. However, our data show that proteases are released into the PVM several hours prior to PVM rupture. Further, the mechanism responsible for PVM poration remains unknown. Thus, it is not clear what prevents premature PVM rupture or how this process is timed. There are several possible scenarios that may explain why PVM poration does not occur until merozoite segmentation is finished. It is possible that the unknown membrane poration protein or complex is not synthesized or released into the PV minutes prior to egress. Another possibility is that the protein responsible for PVM poration is not activated until the PVM rounding occurs, leading to rapid membrane insertion and rupture. A recent study showed that a calcium signal precedes PVM rounding [11] and it is possible that the same signal activates the membrane poration mechanism. Our data suggest that the protein (or complex) responsible for PVM rupture should either be able to form a ~3 μm pore that expands rapidly or this protein (or complex) destabilizes the PVM at a single site that rapidly propagates. Alternatively, it is also possible that a single protein or complex is not responsible for PVM rupture, and that degradation of PV membrane proteins by egress proteases leads to membrane destabilization and rupture, though it is not clear why that would occur at a single site.

The step-by-step pathway that results in intracellular calcium release during egress remains to be determined. Our prior studies identified a key calcium-binding protein, PfERC, that was required for maturation of the egress protease, PMX [20]. However, a later study showed that blocking PMX maturation did not inhibit its proteolytic activity [34]. One possible model that would reconcile these two studies [20,34] would be that PfERC functions in exoneme exocytosis or biogenesis, which could result in inhibition of PMX maturation as well as secretion leading to a block in egress. Therefore, we sought to determine if PfERC functions in exoneme exocytosis or biogenesis. Our data show that knockdown of PfERC results in loss of exoneme exocytosis. These data suggest that PfERC functions upstream of the calcium signal responsible for PVM rounding. Since PfERC is primarily an ER-resident protein, one possibility is that PfERC functions in the biogenesis of exonemes and loss of PfERC leads to defects in biogenesis of exonemes resulting in inhibition of exocytosis. It is likely that defects in exoneme biogenesis may lead to loss of vesicle acidification. However, we did not observe any loss in acidification of exonemes upon PfERC knockdown, these data suggest that any function PfERC may have in exoneme biogenesis does not neutralize exoneme pH. Another possibility is that PfERC may function in an earlier cGMP and/or calcium signal required for exoneme exocytosis. It remains to be determined whether the calcium binding function of PfERC is required for its role in exoneme exocytosis.

These studies demonstrate the development of a live-cell reporter to observe signal-dependent exocytosis required for egress of *P. falciparum* from the infected RBC. Using this reporter, we discovered that signal-dependent exoneme exocytosis starts several hours prior to the egress of merozoites. These data suggest that there may be two independent signals for egress, one for starting exoneme exocytosis several hours prior to egress and another that results in PVM rounding as well as rupture a few minutes before egress (Fig. 7). The data identified a calcium binding protein, PfERC, as a critical player in exoneme exocytosis that may function in the calcium signaling pathway required for exocytosis. These data show that PKG signaling is critical for exoneme exocytosis and occurs several hours prior to egress. It is not yet clear whether this results in a continuous calcium signal or if that occurs later during egress. This study lays the groundwork for dissecting the molecular mechanisms utilized by the deadly malaria-causing parasites to egress from their host cells, which are likely to be similar across the parasitic lifecycle of *P. falciparum*.

**Figure 7:**
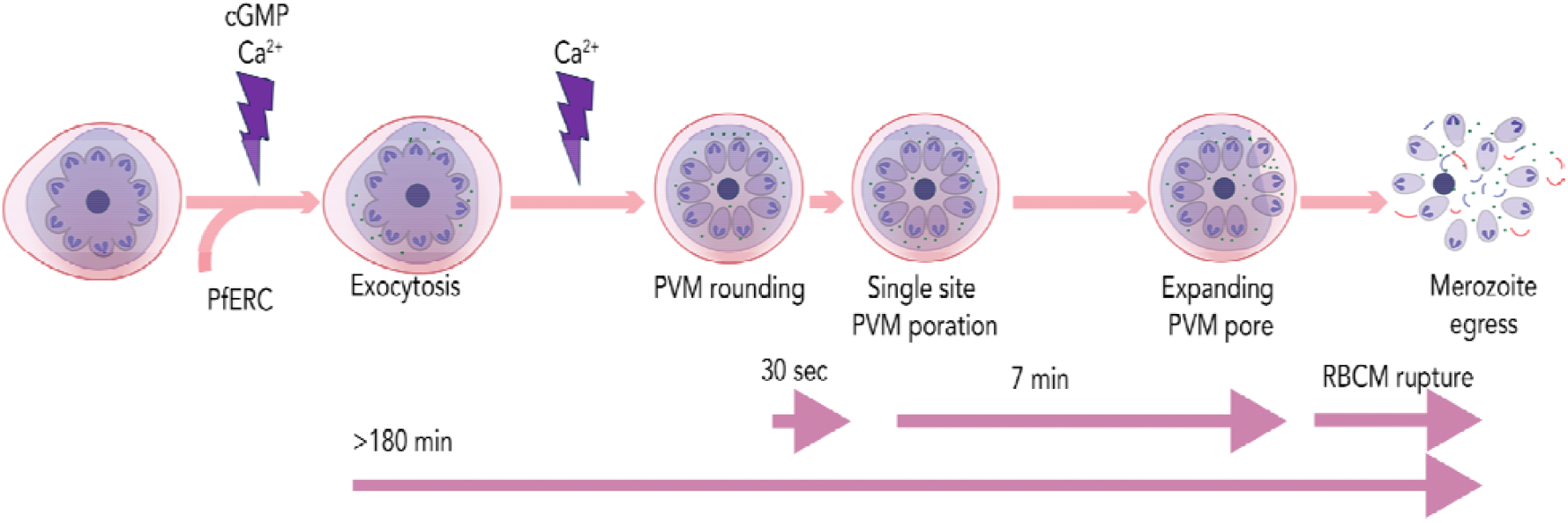
A proposed model for *Plasmodium* egress. Approximately 3 hours prior to egress, an unknown trigger leads to a rise in two second messengers, cGMP and calcium, resulting in the exocytosis of exonemes. This continues until merozoite egress. Exoneme exocytosis also requires PfERC. In 8-10 min prior to egress, a calcium signal triggers the PVM rounding, rapidly followed by its breakdown at a single site. The PVM rupture rapidly expands for minutes before the RBCM ruptures releasing merozoites from the infected RBC.

## Materials and methods

### Plasmid construction

Genomic DNA was isolated from NF54^attb^ *P. falciparum* cultures using a QIAamp DNA Blood Kit (Qiagen), and all constructs were confirmed by sequencing. Briefly, PCR products were inserted into the respective plasmids using the sequence- and ligation-independent cloning (SLIC) method. A mixture of an insert, a cut vector, and T4 DNA polymerase was incubated for 2.5 minutes at room temperature and then for 10 minutes on ice. A bacterial transformation uses a heat shock at 42 °C for 35 sec. All primers utilized in this study are in Table S1. Two plasmids, pbPM2-EXP2-mRuby [29] and pAIO2-EXP2 gRNA [42], were gifts from Dr. Josh Beck, Iowa State University.

SEP was amplified from pHmK-C1 (Addgene) [28] using primers P10/P11. The pPMX-V5-Apt-pMG74 plasmid [20] was modified to replace V5 tag with THE SEP amplicon between PsPXi and AfIII cut site, resulting in a pPMX-SEP-Apt-pMG74 plasmid. A PMX guide RNA targeting the 30 bp upstream *pmx* stop codon was annealed using P12/P13. The annealed fragment was inserted into the plasmid pUF1-Cas9, which contains the DHOD resistance gene, resulting in the plasmid pUF1-Cas9-PMX.

Primers P14/15 were used to amplify an 843 bp fragment of 51 homology PfERC C-terminus (excluding the stop codon) from *P. falciparum* NF54^attb^ genomic DNA. Amplicons were inserted into pHA-SDEL-glmS and pHA-SDEL-M9 using restriction sites SacII and AfeI [20]. A 831-bp 31 homology PfERC 3UTR (beginning at 13 bp downstream PfERC stop codon) was amplified by primers P16/P8 and was inserted into pHA-SDEL-glmS and pHA-SDEL-M9 containing 5 ⍰ homology PfERC C-terminal region at restriction sites HindIII and NheI, resulting in pPfERC-HA-SDEL-glmS or pPfERC-HA-SDEL-M9 [20]. A gRNA targeted at 27 bp upstream of the *erc* stop codon. The PfERC gRNA was annealed using P17/P18 and inserted into the plasmid pUF1-Cas9-guide, resulting in the plasmid pUF1-Cas9-ERC.

### Parasite culturing and transfection

All transfections were done in duplicate. For transfections, an uninfected RBC pellet was electroporated with a matching pair of 20⍰ µg of donor plasmid and 50 µg of CRISPR/Cas9 gRNA plasmid using a Bio-Rad Gene Pulser Xcell Electroporation System. The parasite line was then added to the transfection mixture and cultured in RPMI medium supplemented with Albumax I (Gibco) and the required drugs described previously [20,43–46]. All primers used are in Table S1.

To generate PfEXP2^mRuby3^ parasites, 20 ⍰ μg of the pbPM2-EXP2-mRuby donor plasmid was linearized with AflII overnight, and 50 μg of the pAIO-EXP2-gRNA were co-transfected into NF54^attb^ parasites [47]. At 24 h after transfection, 2.5⍰ μg/ml of Blasticidin-S (BSD) was added to select for pbPM2-EXP2-mRuby expression [29,39]. The parasite population was cloned by limiting dilution. A PCR diagnostic for integration was performed using primers P1/P3 to amplify a 2.1 kb fragment corresponding to the correct integrant.

For generating PfEXP2^mRuby3^/PMX^SEP^ parasites, 20 1μg of pPMX-SEP-Apt-pMG74, a donor plasmid, was linearized with EcoRV overnight, and 50 μg of pUF1-Cas9-PMX gRNA plasmid were co-transfected into PfEXP2^mRuby3^ parasites [13,20]. Transfected parasites were grown in 0.5 μM anhydrous tetracycline (aTc) (Cayman Chemical) and 2.5⍰ μg/ml of Blasticidin-S (BSD) [20,39]. At 48 h post-transfection, Cas9 expression was selected by drug cycling [48,49] in the presence and absence of 11μM DSM1 for 4 days until parasites were detected in the culture. Next, the parasite population was cloned by limiting dilution. A PCR diagnostic for integration was performed using primers P4/P6 to amplify a 2.8 kb fragment corresponding to the correct integrant.

For generating the PfERC-glmS and PfERC-M9 parasites, a mix of two plasmids (20 μg of a pPfERC-HA-SDEL-glmS or pPfERC-HA-SDEL-M9 donor plasmid and 50 μg of pUF1-Cas9-ERC guide plasmid) was co-transfected into NF54^attb^ parasites. To select for Cas9 expression,11μM DSM1 was applied to the transfected culture as previously described [20]. Clonal parasites were subjected to PCR diagnosis using primers P7/P8 and P7/P9 to amplify a 2.2-kb and 1.1-kb fragment, respectively, for correct integration [20].

To generate PfERC-*glmS*/PMX^SEP^ and PfERC-*M9*/PMX^SEP^ parasites, 20 1μg of pPMX-SEP-Apt-pMG74, a donor plasmid, was linearized with EcoRV overnight, and a 50 μg of pUF1-Cas9-PMX gRNA plasmid were co-transfected into PfERC-glmS or PfERC-M9 parasites [13,20]. Transfected cultures were grown in 2.5 μg/ml of blasticidin S (BSD) [20,39]. To select a donor plasmid, 0.5 μM anhydrous tetracycline (aTc) (Cayman Chemical) was applied at 48 h post-transfection. A PCR diagnostic for integration was performed using primers P4/P6 to amplify a 2.8 kb fragment corresponding to the correct integrant.

### Western blotting

Western blotting was performed using *P. falciparum* lysates as described previously [20] with minor modifications. Highly synchronized late-schizont parasite culture pellets were treated with ice-cold 0.05% saponin–phosphate-buffered saline (PBS) for 15min, followed by sonication. The sonicated lysates were collected for detection of proteins separated via a western blot assay. The antibodies used in this study were rabbit anti-HA (715500; Invitrogen) (1:1000), mouse anti-GFP (JL-8; Takara) (1:3000), rabbit anti-EXP2 (from H.Ke) (1:10000), and mouse anti-GAPDH (1.4; Millipore Sigma; 1:2000). The secondary antibodies used were IRDye 680CW goat anti-rabbit IgG and IRDye 800CW goat anti-mouse IgG (Li-COR Biosciences) (1:20,000). The Western blotting images were processed using Odyssey Clx Li-COR infrared imaging system software (Li-COR Biosciences).

### Time-lapse imaging of live P. falciparum

Late-stage schizont parasites (44 h.p.i) were isolated on a Percoll gradient (Genesee Scientific). Percoll pellet was incubated in the PKG inhibitors either 1.5 μM compound 1 (C1) {4-[2-(4-fluorophenyl)-5-(1-methylpiperidine-4-yl)-1H-pyrrol-3-yl] pyridine} [8,18] or 25 nM ML10 (BEI Resources, catalog no. NR-56525, www.beiresources.org) [50] at 37 °C in a CO2 incubator. After 4 h incubation, the egress inhibitor was washed off twice using pre-warmed 37°C cRPMI media. After washing off the egress inhibitor, 200-500 μl schizont-stage parasite culture was spun down at 3000 rpm for 30-sec, and the supernatant was removed. The pellet was gently resuspended with warm (37°C) RMPI media, and the tube was spun down at 3000 rpm for 1 min. After removing the supernatant, the pellet was gently resuspended with 200-500 μl warm (37°C) media before being placed onto a 35 mm glass-bottom dish coated with poly-L-lysine for imaging.

The 30-minute time-lapse video was performed using a DeltaVision II microscope with autofocus. The PfEXP2^mRuby3^/PMX^SEP^ schizonts were imaged with a Live cell filter set CFP, YFP, GFP, and mCherry. To minimize photodamage, the laser power level was kept at 10% (or less). A 488-nm-wavelength laser was used to excite PMX^SEP^, and a 561-nm-wavelength laser was used to excite PfEXP2^mRuby3^. Time-lapse images were taken every 30 sec up to 30 min at 37 °C in an imaging chamber supplemented with CO2.

The 5-h time-lapse video was imaged using a Zeiss LSM 980 Confocal system using an inverted Axio Observer 7 microscope stand with transmitted (HAL), UV (HBO), and laser (405 – 730 nm) illumination sources. A 488-nm-wavelength laser (10% power) was used to excite PMX^SEP^, and a 561-nm-wavelength laser (2% power) was used to excite PfEXP2^mRuby3^. The microscope system uses Zen 3.3 (Blue) acquisition software with autofocus. The dark incubation chamber with regulated CO2 (5%), temperature (37 °C), and humidity was used. Images were captured every 5 minutes for 5 hours. To treat parasites with BaF, highly synchronized PfEXP2^mRuby3^/PMX^SEP^ parasites were prepared as described. The 44 h.p.i schizont pellet was incubated with ML10 for 2-4 h. Before imaging, 100 nM BaF (Cayman Chemical) was added to the culture and incubated for 10 min.

### Image processing and analysis

ImageJ (Fiji version 2.0.0-rc-68/1.52h) was used to crop images, adjust brightness and intensity, overlay channels, and prepare montages. The adjustments to brightness and contrast were made only for display purposes. From the Analyze menu, set measurements were selected with the area, min, max gray values, and mean gray values. The cell of interest was manually selected using a freeform tool, and this step was repeated for every time point. The measure menu was used to measure fluorescence. The fluorescence intensity of each cell was calculated using the calculation for corrected total cell fluorescence (CTCF) = integrated density–(area of selected cell × mean fluorescence of background readings) [51,52]. Plots and statistical analysis (2-sided unpaired Student t-tests) were done using GraphPad Prism 10.

## Acknowledgements

We thank members of the Muralidharan lab and Matthias Garten for their thoughtful comments and suggestions; Josh Beck for the EXP2 tagging plasmids; Hangjun Ke for anti-EXP2 antibodies; Purnima Bhanot for compound 1; Muthugapatti Kandasamy at the University of Georgia (UGA) Biomedical Microscopy Core. This work was supported by grants from the U.S. National Institutes of Health to V.M. (R56AI173133 and R01AI179950) and M.A.F. (T32AI060546).

## Supplementary Data

**Supplementary Figure 1.**
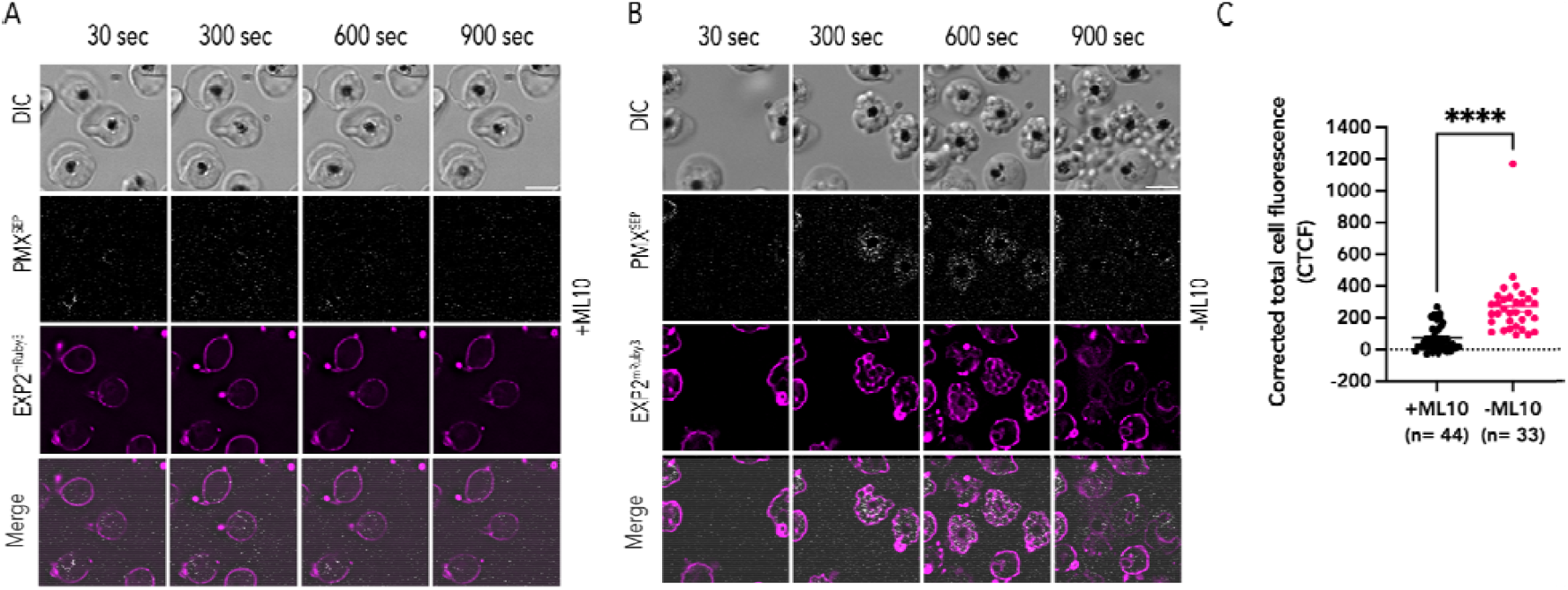
ML10 PKG inhibition blocks exoneme exocytosis. **(A)** A representative image from live imaging of synchronized PfEXP2^mRuby3^/PMX^SEP^ schizonts in the presence of ML10 (**B**) A representative image from live imaging of synchronized PfEXP2^mRuby3^/PMX^SEP^ schizonts after ML10 removal. Parasite egress occurs, and free-merozoites are scattered in the extracellular space (DIC). **(C)** Corrected total cell fluorescence (CTCF) of PMX^SEP^ quantified from time-lapse images of synchronized PfEXP2^mRuby3^/PMX^SEP^ schizonts incubated with ML 10 (black; n=44, 2 biological replicates) or without ML10 (magenta; n=33, 2 biological replicates) C1. ***p-value<0.0001, unpaired t-test. error bars = SEM Scale bar=5⍰μm.

**Supplementary Figure 2:**
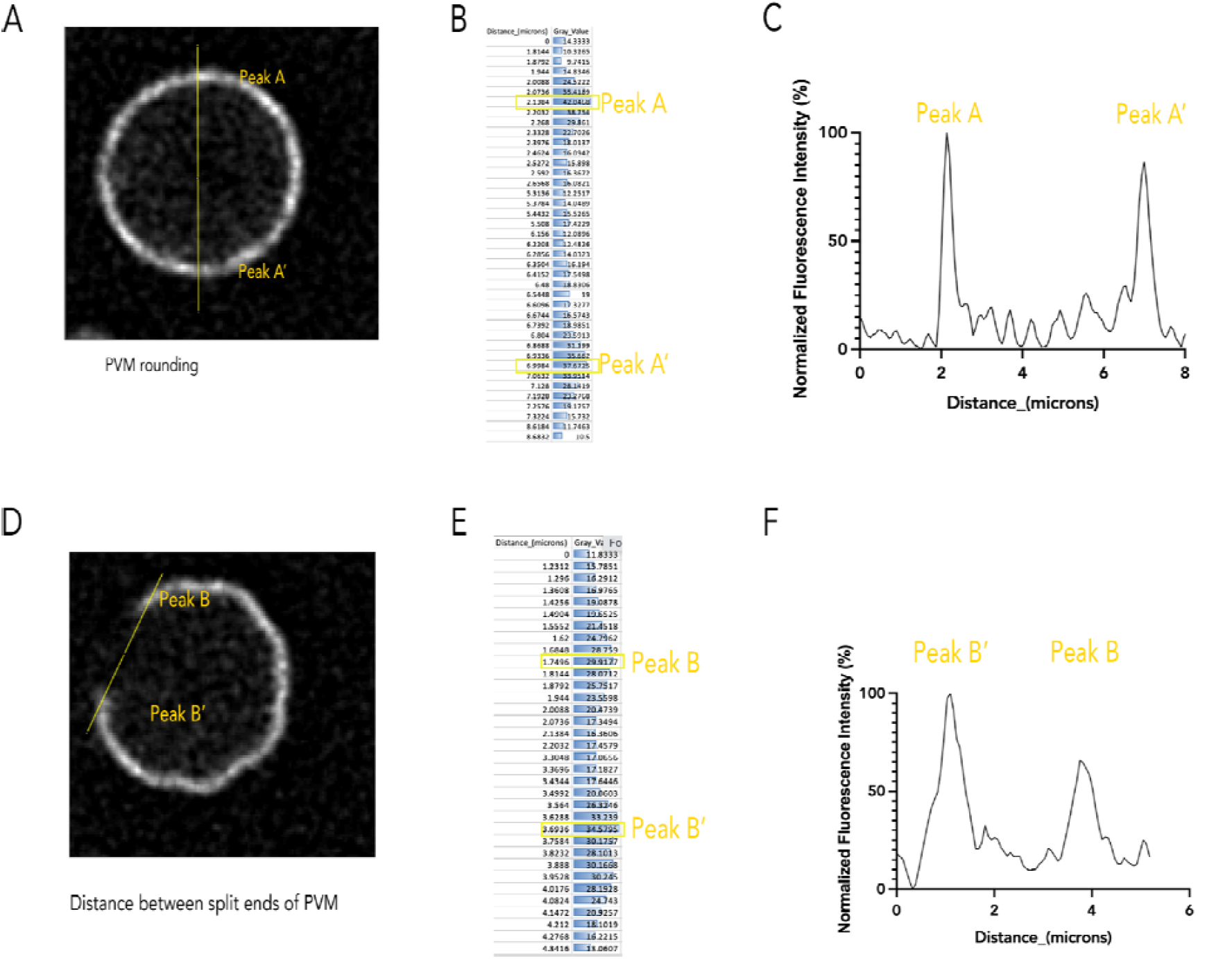
Measuring PVM diameter and split ends. A representative of measuring PVM diameter (A-C). By drawing a line across the rounded PVM (A), measuring fluorescence intensity values corresponding to the lines are shown in (B), and generating a normalized fluorescence intensity graph in (C), the fluorescence intensity peaks are labelled as A and A’. A representative of measuring the distance between two split ends of PVM (D-F). Drawing a line across two split ends labeled as Peak B and B’ (C) and measuring fluorescence intensity values corresponding to the line (D) and its normalized fluorescence intensity graph generated (F).

**Supplementary Figure 3:**
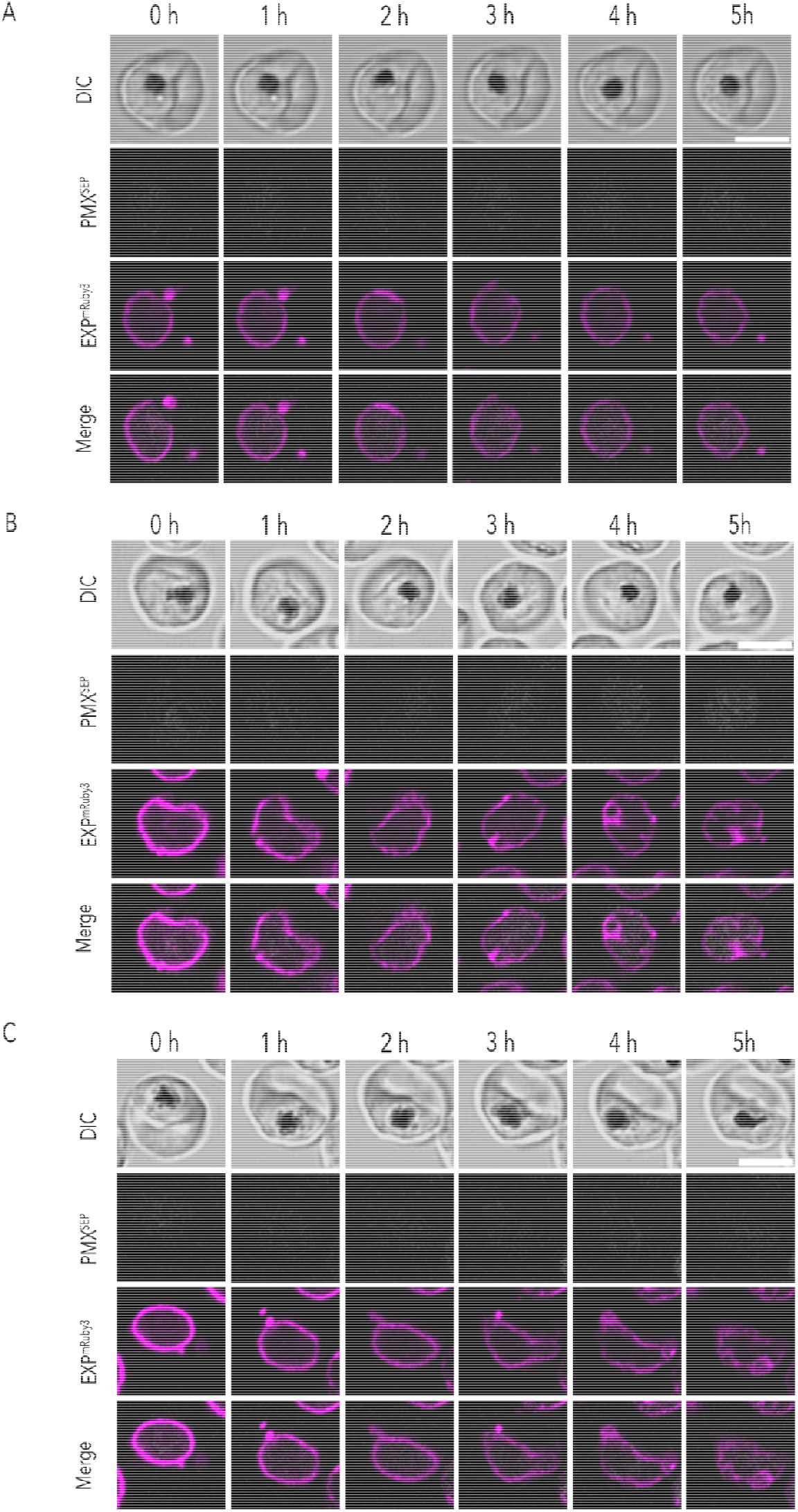
Exonemes are acidified prior to exocytosis. Three representative images of live cell imaging of PfEXP2^mRuby3^/PMX^SEP^ schizonts in the presence of ML10 (A-C). All schizonts show no fluorescence detected throughout the 5-hour recording. Time-lapse images were taken at 5-min intervals. Scale bar=5 μm; n= 2 biological replicates.

**Supplementary Figure 4:**
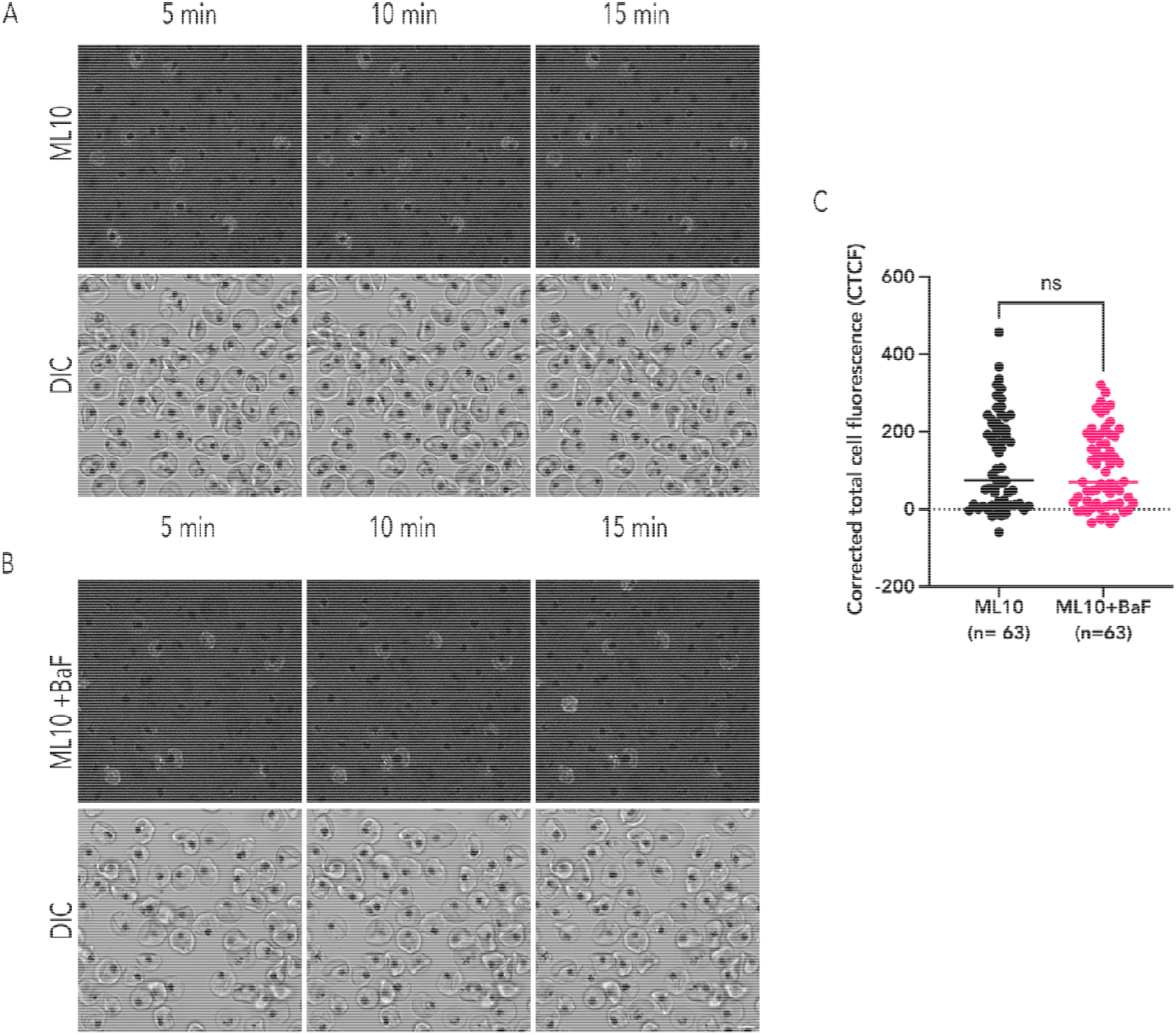
Exoneme acidification is insensitive to V-type ATPases-specific inhibitor. (A) A representative still image from a time-lapse video of ML10-treated PfEXP2^mRuby3^/PMX^SEP^ late schizont and (B) A representative still image from a time-lapse video of BaF-ML10-treated PfEXP2^mRuby3^/PMX^SEP^ late schizont. (C) Corrected total cell fluorescence (CTCF) values of PMX^SEP^ schizonts. Black dot represents the CTCF value of each schizont with ML10 (n=63, 3 replicates). Pink dot represents the CTCF value of each schizont in the presence of ML10 and BaF (n= 63, 3 replicates), ns= not significant, unpaired t-test Scale bar=5⍰μm.

**Supplementary Figure 5:**
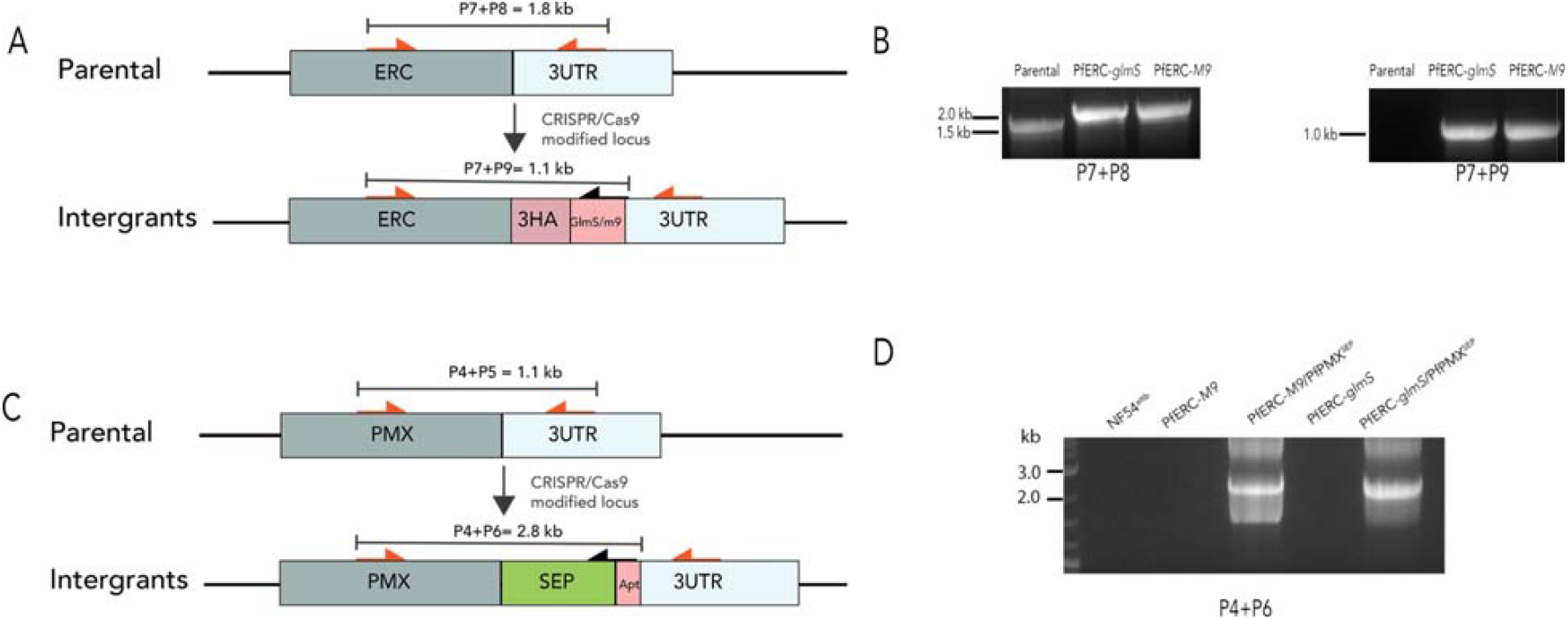
Generation of PfERC conditional mutants expressing PMX^SEP^. **(A)** Schematics of the targeting (pERC-TOPO) plasmids by the CRISPR/Cas9 system and a guide RNA. The locations of diagnostic primers (P7, P9, and P9) used to demonstrate the repair of the locus via double-crossover homologous integration are shown. (B) Agarose gel showing the PCR diagnostic test using the primer pairs P7+P8 (left) and P7+P9 (right). Amplicons were amplified from the genomic DNA extracted from parental and mutant parasites. **(C)** Schematics of the fusion of *pmx* with SEP. (D) PCR diagnosis of SEP into the PfPMX locus in NF54^attb^, PfERC-*M9*, PfERC-*M9*/PMX^SEP^, PfERC-*glmS* and PfERC-*glmS*/PMX^SEP^ using primer pair (P4 and P6).

**Table S1.**
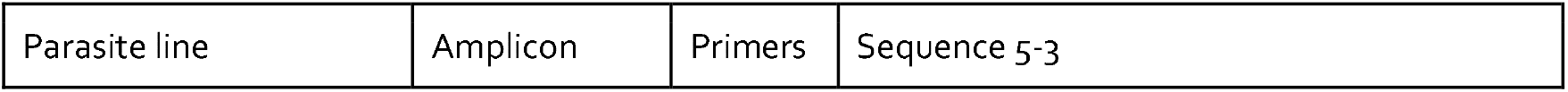
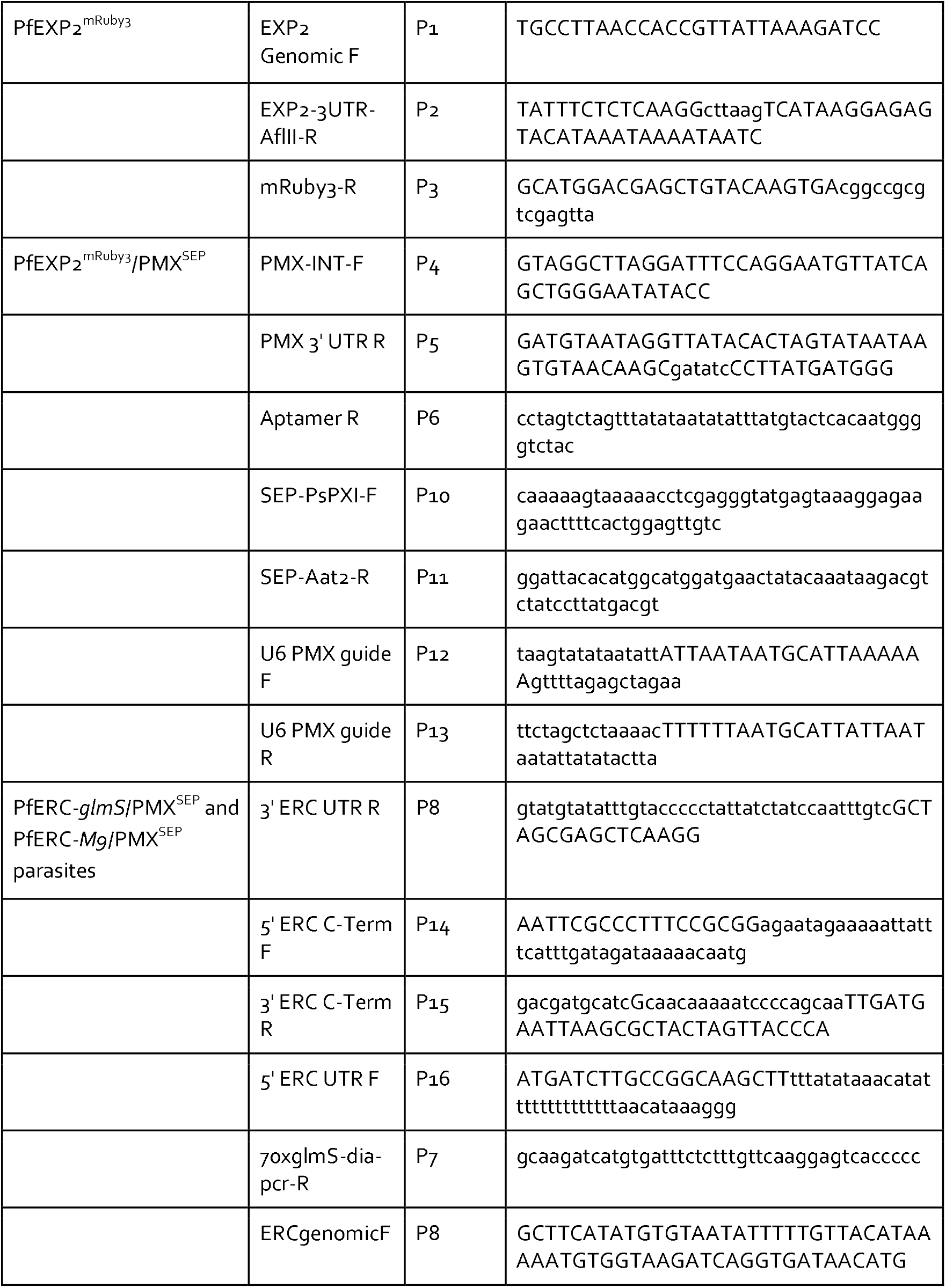
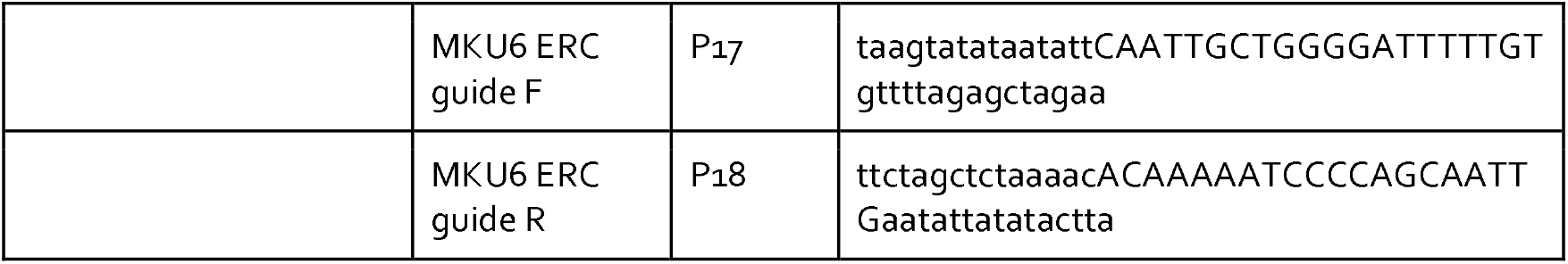
List of primers used in the study to generate the parasite lines.

**Movie S1: Time-lapse imaging of PfEXP2**^**mRuby3**^**/PMX**^**SEP**^ **schizonts in the presence of C1**. Representatives of PfEXP2^mRuby3^/PMX^SEP^ parasites within RBCs until the end of recording (DIC, left). PfEXP2^mRuby3^ signals were irregularly shaped, and PMX^SEP^ was not detected (Merged two fluorescence channels, right) (5 recordings from 3 biological replicates). Scale bar=51μm

**Movie S2: Time-lapse imaging of PfEXP2**^**mRuby3**^**/PMX**^**SEP**^ **schizonts in the absence of C1**. Representative of PfEXP2^mRuby3^/PMX^SEP^ parasites egress upon C1 removal, and free merozoites into the extracellular space (left). PMX^SEP^ fluorescence was detected from the beginning of the recording until after merozoite egress occured. PfEXP2^mRuby3^ signals were irregularly shaped (right), (n = 6 recordings from 3 biological replicates). Scale bar=5⍰μm

**Movie S3: Time-lapse imaging of exocytosis and egress of PfEXP2**^**mRuby3**^**/PMX**^**SEP**^ **schizonts**. Late schizont PfEXP2^mRuby3^/PMX^SEP^ parasites (left). Exoneme exocytosis of PfEXP2^mRuby3^/PMX^SEP^ was detected as punctate patterns (Middle). PVM morphology changed from irregular shape to a rounded shape before PVM breakdown (right), (7 biological replicates, double-labeled parasites, time-lapse recordings of parasite egress are at a 30-sec interval). Scale bar=5⍰μm

**Movie S4: Time-lapse imaging of PfEXP2**^**mRuby3**^**/PMX**^**SEP**^ **schizonts at 4 sec-intervals shows PVM rupture at a single site**. Representative of still images from 4-sec intervals of PfEXP2^mRuby3^/PMX^SEP^ late schizont (left). PfEXP2^mRuby3^ signals were irregularly shaped and transitioned to rounded before rupture at a single site (right). Scale bar=5⍰μm (5 recordings from 2 biological replicates)

**Movie S5: Time-lapse imaging of PfEXP2**^**mRuby3**^**/PMX**^**SEP**^ **schizonts at 2 sec-intervals showed the PVM rupture at a single site**. Representative of PfEXP2^mRuby3^ images showing PVM morphology changed from irregular to rounded before its breakdown at a single site. Scale bar=5 μm (6 recordings from 4 biological replicates)

**Movie S6: Exocytosis of PfEXP2**^**mRuby3**^**/PMX**^**SEP**^ **schizonts was detected over 3 hrs before egress**. Punctate PMX^SEP^ exocytosis signal was detected about 3 hours prior to egress (left). PfEXP2^mRuby3^ signals showed PVM morphology (middle). PfEXP2^mRuby3^/PMX^SEP^ schizont egressed freeing merozoites into the extracellular space (right). Scale bar=5⍰μm

**Movie S7: Exonemes are acidified at least 5 hours prior to egress**. PMX^SEP^ fluorescence was not detected in the presence of ML10 throughout 5 hours of recording (right) showing that exonemes were already acidified several hours prior to egress. As expected PfEXP2^mRuby3^/PMX^SEP^ schizonts could not egress in the presence of ML10 (left). Time-lapse images were taken at 5-min intervals. Scale bar=5⍰μm; n= 2 biological replicates.

**Movie S8: Time-lapse imaging of GlcN-treated PfERC-**M9**/PMX**^**SEP**^ **schizonts**. Representatives of GlcN-treated PfERC-*M9*/PMX^SEP^ schizonts egress upon C1 removal (left). PMX^SEP^ fluorescence signal is detected in schizonts indicating exoneme exocytosis (right; 10 recordings, n=3 biological replicates). Scale bar=5⍰μm

**Movie S9: Glc-treated PfERC-**glmS**/PMX**^**SEP**^ **schizonts fail to egress**. Representatives of GlcN-treated PfERC-*glmS*/PMX^SEP^schizonts egress upon C1 removal reside within the red blood cells (left). Almost all GlcN-treated PfERC-*glmS*/PMX^SEP^ schizonts lack PMX^SEP^ fluorescence indicating loss of exoneme exocytosis (right; 11 recordings, n=3 biological replicates). Scale bar=5⍰μm

## REFERENCES

1. World Health Organization 2024. [cited 4 Aug 2025]. Available: https://www.who.int/teams/global-malaria-programme/reports/world-malaria-report-2024

2. Tan MSY, Blackman MJ. Malaria parasite egress at a glance. J Cell Sci. 2021;134.

3. Dvorin JD, Martyn DC, Patel SD, Grimley JS, Collins CR, Hopp CS, et al. A plant-like kinase in Plasmodium falciparum regulates parasite egress from erythrocytes. Science. 2010;328: 910–912.

4. Singh S, Alam MM, Pal-Bhowmick I, Brzostowski JA, Chitnis CE. Distinct external signals trigger sequential release of apical organelles during erythrocyte invasion by malaria parasites. PLoS Pathog. 2010;6: e1000746.

5. Farrell A, Thirugnanam S, Lorestani A, Dvorin JD, Eidell KP, Ferguson DJP, et al. A DOC2 protein identified by mutational profiling is essential for apicomplexan parasite exocytosis. Science. 2012;335: 218–221.

6. Agarwal S, Singh MK, Garg S, Chitnis CE, Singh S. Ca(2+)-mediated exocytosis of subtilisin-like protease 1: a key step in egress of Plasmodium falciparum merozoites. Cell Microbiol. 2013;15: 910–921.

7. Gao X, Gunalan K, Yap SSL, Preiser PR. Triggers of key calcium signals during erythrocyte invasion by Plasmodium falciparum. Nat Commun. 2013;4: 2862.

8. Collins CR, Hackett F, Strath M, Penzo M, Withers-Martinez C, Baker DA, et al. Malaria parasite cGMP-dependent protein kinase regulates blood stage merozoite secretory organelle discharge and egress. PLoS Pathog. 2013;9: e1003344.

9. Brochet M, Collins MO, Smith TK, Thompson E, Sebastian S, Volkmann K, et al. Phosphoinositide metabolism links cGMP-dependent protein kinase G to essential Ca2+ signals at key decision points in the life cycle of malaria parasites. PLoS Biol. 2014;12: e1001806.

10. Yeoh S, O’Donnell RA, Koussis K, Dluzewski AR, Ansell KH, Osborne SA, et al. Subcellular discharge of a serine protease mediates release of invasive malaria parasites from host erythrocytes. Cell. 2007;131: 1072–1083.

11. Seveno M, Loubens MN, Berry L, Graindorge A, Lebrun M, Lavazec C, et al. The malaria parasite PP1 phosphatase controls the initiation of the egress pathway of asexual blood-stages by regulating the rounding-up of the vacuole. PLoS Pathog. 2025;21: e1012455.

12. Dvorin JD, Goldberg DE. Plasmodium Egress Across the Parasite Life Cycle. Annu Rev Microbiol. 2022;76: 67–90.

13. Nasamu AS, Glushakova S, Russo I, Vaupel B, Oksman A, Kim AS, et al. Plasmepsins IX and X are essential and druggable mediators of malaria parasite egress and invasion. Science. 2017;358: 518–522.

14. Pino P, Caldelari R, Mukherjee B, Vahokoski J, Klages N, Maco B, et al. A multistage antimalarial targets the plasmepsins IX and X essential for invasion and egress. Science. 2017;358: 522–528.

15. Favuzza P, de Lera Ruiz M, Thompson JK, Triglia T, Ngo A, Steel RWJ, et al. Dual Plasmepsin-Targeting Antimalarial Agents Disrupt Multiple Stages of the Malaria Parasite Life Cycle. Cell Host Microbe. 2020;27: 642–658.e12.

16. Thomas JA, Tan MSY, Bisson C, Borg A, Umrekar TR, Hackett F, et al. A protease cascade regulates release of the human malaria parasite Plasmodium falciparum from host red blood cells. Nat Microbiol. 2018;3: 447–455.

17. Ruecker A, Shea M, Hackett F, Suarez C, Hirst EMA, Milutinovic K, et al. Proteolytic activation of the essential parasitophorous vacuole cysteine protease SERA6 accompanies malaria parasite egress from its host erythrocyte. J Biol Chem. 2012;287: 37949–37963.

18. Hale VL, Watermeyer JM, Hackett F, Vizcay-Barrena G, van Ooij C, Thomas JA, et al. Parasitophorous vacuole poration precedes its rupture and rapid host erythrocyte cytoskeleton collapse in Plasmodium falciparum egress. Proceedings of the National Academy of Sciences. 2017;114: 3439–3444.

19. Glushakova S, Beck JR, Garten M, Busse BL, Nasamu AS, Tenkova-Heuser T, et al. Rounding precedes rupture and breakdown of vacuolar membranes minutes before malaria parasite egress from erythrocytes. Cell Microbiol. 2018;20: e12868.

20. Fierro MA, Asady B, Brooks CF, Cobb DW, Villegas A, Moreno SNJ, et al. An Endoplasmic Reticulum CREC Family Protein Regulates the Egress Proteolytic Cascade in Malaria Parasites. MBio. 2020;11: e03078–19

21. Honoré B. The rapidly expanding CREC protein family: members, localization, function, and role in disease. Bioessays. 2009;31: 262–277.

22. Lam PPL, Hyvärinen K, Kauppi M, Cosen-Binker L, Laitinen S, Keränen S, et al. A cytosolic splice variant of Cab45 interacts with Munc18b and impacts on amylase secretion by pancreatic acini. Mol Biol Cell. 2007;18: 2473–2480.

23. Miesenböck G, De Angelis DA, Rothman JE. Visualizing secretion and synaptic transmission with pH-sensitive green fluorescent proteins. Nature. 1998;394: 192–195.

24. Sankaranarayanan S, De Angelis D, Rothman JE, Ryan TA. The use of pHluorins for optical measurements of presynaptic activity. Biophys J. 2000;79: 2199–2208.

25. Li S-A, Meng X-Y, Zhang Y-J, Chen C-L, Jiao Y-X, Zhu Y-Q, et al. Progress in pH-Sensitive sensors: essential tools for organelle pH detection, spotlighting mitochondrion and diverse applications. Front Pharmacol. 2023;14: 1339518.

26. Alder A, Sanchez CP, Russell MRG, Collinson LM, Lanzer M, Blackman MJ, et al. The role of Plasmodium V-ATPase in vacuolar physiology and antimalarial drug uptake. Proc Natl Acad Sci U S A. 2023;120: e2306420120.

27. Martineau M, Somasundaram A, Grimm JB, Gruber TD, Choquet D, Taraska JW, et al. Semisynthetic fluorescent pH sensors for imaging exocytosis and endocytosis. Nat Commun. 2017;8: 1412.

28. Tanida I, Ueno T, Uchiyama Y. A super-ecliptic, pHluorin-mKate2, tandem fluorescent protein-tagged human LC3 for the monitoring of mammalian autophagy. PLoS One. 2014;9: e110600.

29. Garten M, Beck JR, Roth R, Tenkova-Heuser T, Heuser J, Istvan ES, et al. Contacting domains segregate a lipid transporter from a solute transporter in the malarial host-parasite interface. Nat Commun. 2020;11: 3825.

30. Baker DA, Stewart LB, Large JM, Bowyer PW, Ansell KH, Jiménez-Díaz MB, et al. A potent series targeting the malarial cGMP-dependent protein kinase clears infection and blocks transmission. Nat Commun. 2017;8: 430.

31. Shadija N, Dass S, Xu W, Ke H. Multifunctionality of V-type ATPase during asexual growth and development of Plasmodium falciparum. bioRxiv. 2023. p. 2023.08.02.551680. doi:10.1101/2023.08.02.551680

32. Prommana P, Uthaipibull C, Wongsombat C, Kamchonwongpaisan S, Yuthavong Y, Knuepfer E, et al. Inducible knockdown of Plasmodium gene expression using the glmS ribozyme. PLoS One. 2013;8: e73783.

33. Ganesan SM, Falla A, Goldfless SJ, Nasamu AS, Niles JC. Synthetic RNA–protein modules integrated with native translation mechanisms to control gene expression in malaria parasites. Nat Commun. 2016;7: 10727.

34. Mukherjee S, Nguyen S, Sharma E, Goldberg DE. Maturation and substrate processing topography of the Plasmodium falciparum invasion/egress protease plasmepsin X. Nat Commun. 2022;13: 4537.

35. Mukherjee S, Nasamu AS, Rubiano KC, Goldberg DE. Activation of the Plasmodium Egress Effector Subtilisin-Like Protease 1 Is Mediated by Plasmepsin X Destruction of the Prodomain. MBio. 2023;14: e0067323.

36. Martinez M, Bouillon A, Brûlé S, Raynal B, Haouz A, Alzari PM, et al. Prodomain-driven enzyme dimerization: a pH-dependent autoinhibition mechanism that controls Plasmodium Sub1 activity before merozoite egress. MBio. 2024; e0019824.

37. Withers-Martinez C, Strath M, Hackett F, Haire LF, Howell SA, Walker PA, et al. The malaria parasite egress protease SUB1 is a calcium-dependent redox switch subtilisin. Nat Commun. 2014;5: 3726.

38. Solebo O, Ling L, Nwankwo I, Zhou J, Fu T-M, Ke H. Plasmodium falciparum utilizes pyrophosphate to fuel an essential proton pump in the ring stage and the transition to trophozoite stage. PLoS Pathog. 2023;19: e1011818.

39. Ganesan SM, Morrisey JM, Ke H, Painter HJ, Laroiya K, Phillips MA, et al. Yeast dihydroorotate dehydrogenase as a new selectable marker for Plasmodium falciparum transfection. Mol Biochem Parasitol. 2011;177: 29–34.

40. Rudlaff RM, Kraemer S, Streva VA, Dvorin JD. An essential contractile ring protein controls cell division in Plasmodium falciparum. Nat Commun. 2019;10: 2181.

41. Tan MSY, Koussis K, Withers-Martinez C, Howell SA, Thomas JA, Hackett F, et al. Autocatalytic activation of a malarial egress protease is druggable and requires a protein cofactor. EMBO J. 2021;40: e107226.

42. Glushakova S, Busse BL, Garten M, Beck JR, Fairhurst RM, Goldberg DE, et al. Exploitation of a newly-identified entry pathway into the malaria parasite-infected erythrocyte to inhibit parasite egress. Sci Rep. 2017;7: 1–13.

43. Drew ME, Banerjee R, Uffman EW, Gilbertson S, Rosenthal PJ, Goldberg DE. Plasmodium food vacuole plasmepsins are activated by falcipains. J Biol Chem. 2008;283: 12870–12876.

44. Florentin A, Stephens DR, Brooks CF, Baptista RP, Muralidharan V. Plastid biogenesis in malaria parasites requires the interactions and catalytic activity of the Clp proteolytic system. Proc Natl Acad Sci U S A. 2020;117: 13719–13729.

45. Florentin A, Cobb DW, Fishburn JD, Cipriano MJ, Kim PS, Fierro MA, et al. PfClpC Is an Essential Clp Chaperone Required for Plastid Integrity and Clp Protease Stability in Plasmodium falciparum. Cell Rep. 2017;21: 1746–1756.

46. Cobb DW, Florentin A, Fierro MA, Krakowiak M, Moore JM, Muralidharan V. The Exported Chaperone PfHsp70x Is Dispensable for the Plasmodium falciparum Intraerythrocytic Life Cycle. mSphere. 2017;2: e00363–17

47. Nkrumah LJ, Muhle RA, Moura PA, Ghosh P, Hatfull GF, Jacobs WR Jr, et al. Efficient site-specific integration in Plasmodium falciparum chromosomes mediated by mycobacteriophage Bxb1 integrase. Nat Methods. 2006;3: 615–621.

48. Harris PK, Yeoh S, Dluzewski AR, O’Donnell RA, Withers-Martinez C, Hackett F, et al. Molecular identification of a malaria merozoite surface sheddase. PLoS Pathog. 2005;1: 241–251.

49. Collins CR, Das S, Wong EH, Andenmatten N, Stallmach R, Hackett F, et al. Robust inducible Cre recombinase activity in the human malaria parasite Plasmodium falciparum enables efficient gene deletion within a single asexual erythrocytic growth cycle. Mol Microbiol. 2013;88: 687–701.

50. Ressurreição M, Thomas JA, Nofal SD, Flueck C, Moon RW, Baker DA, et al. Use of a highly specific kinase inhibitor for rapid, simple and precise synchronization of Plasmodium falciparum and Plasmodium knowlesi asexual blood-stage parasites. PLoS One. 2020;15: e0235798.

51. McCloy RA, Rogers S, Caldon CE, Lorca T, Castro A, Burgess A. Partial inhibition of Cdk1 in G 2 phase overrides the SAC and decouples mitotic events. Cell Cycle. 2014;13: 1400–1412.

52. Jakic B, Buszko M, Cappellano G, Wick G. Elevated sodium leads to the increased expression of HSP60 and induces apoptosis in HUVECs. PLoS One. 2017;12: e0179383.

